# Water wisteria genome reveals environmental adaptation and heterophylly regulation in amphibious plants

**DOI:** 10.1101/2022.09.19.508473

**Authors:** Gaojie Li, Xuyao Zhao, Jingjing Yang, Shiqi Hu, Jathish Ponnu, Seisuke Kimura, Inhwan Hwang, Keiko U Torii, Hongwei Hou

## Abstract

Heterophylly is a phenomenon in which an individual plant dramatically changes its leaf shape in response to the surrounding environment. *Hygrophila difformis* (Acanthaceae), also known as water wisteria, has recently emerged as a model plant to study heterophylly because of its striking leaf shape variation in response to various ecological factors. Under submerged conditions, *H. difformis* develops complex leaves and in terrestrial conditions it develops simple leaves. Here, we sequenced and assembled the chromosome-level genome of triploid *H. difformis* (scaffold N50: 60.43 Mb, genome size: 871.92 Mb), which reveals 36,099 predicted protein-coding genes distributed over 15 pseudochromosomes. *H. difformis* diverged from its relatives during the Oligocene climate-change period and expanded the gene families related to its amphibious lifestyle. Genes involved in environmental stimuli, leaf development, and other pathways are differentially expressed in submerged and terrestrial conditions, possibly modulating morphological and physiological acclimation to changing environments. We confirmed that auxin plays a role in the heterophylly of *H. difformis*. Finally, we discovered candidate genes that respond to different environmental conditions and elucidated the role of *LATE MERISTEM IDENTITY 1* (*LMI1*) in heterophylly. Our study establishes *H. difformis* as a model for studying the interconnections between ecological adaptation and plant morphological features.

## Introduction

Amphibious plants can grow both in water and on land, switching their morphology and physiology according to various environmental cues (Agarie et al., 1997). While genomic data are available for several aquatic plants (Lan et al., 2017; Xue et al., 2020; Lu et al., 2021; Abramson et al., 2022; Shi et al., 2022), amphibious plants remain largely unexplored at the genomic level. Climate change poses new challenges for plant growth and survival. One of the strategies that plants use to cope with these challenges is phenotypic plasticity (Alpert and Simms, 2002), the ability to change their phenotypes in response to environmental cues. Phenotypic plasticity connects plant evolution, ecology and molecular biology and helps to predict plant responses to novel environments under climate change (Nicotra et al., 2010; Arnold et al., 2019; Stotz et al., 2021). A striking example of phenotypic plasticity is heterophylly, the ability of a plant to alter its leaf shape depending on the environment (van Veen and Sasidharan, 2021). Heterophylly is common in amphibious plants, which can grow both in land and water (Nakayama et al., 2014; Li et al., 2017; Kim et al., 2018; Koga et al., 2021). Heterophyllous plants can adjust their leaf morphology to various environmental factors such as submergence (Li et(Han et al., 2021). al., 2017), light (Momokawa et al., 2011; Nakayama et al., 2014), temperature (Nakayama and Kimura, 2015), and CO_2_ (Titus and Gary Sullivan, 2001), bridging plant ecology, evolution, and molecular biology, might enable to predict plant responses to novel environments posed by climate change. Whereas studies on model plants, like *Arabidopsis thaliana*, have greatly contributed to identifying genes and processes regulating plant phenotypes, we need a suitable model system on phenotypic plasticity to understand the relationships of environmental factors on plant phenotypes. Interestingly, heterophyllous plants across diverse taxa often show similar leaf variations to adapt to the amphibious environment, making them potential plant models for convergent evolution and environmental adaptation.

Recently, several mechanisms regulating heterophylly have been suggested in a few amphibious species. For example, ethylene and abscisic acid (ABA) control genes that regulate adaxial–abaxial leaf polarity and induce heterophylly in *Ranunculus trichophyllus* (Ranunculaceae) (Kim et al., 2018). Similarly, genes that affect cell elongation and division contribute to leaf differentiation in *Callitriche palustris* (Plantaginaceae) (Koga et al., 2021). In *Rorippa aquatica* (Brassicaceae), *KNOTTED1-LIKE HOMEOBOX* (*KNOX1*) genes modulate the levels of endogenous phytohormones gibberellic acid (GA) and cytokinin to influence temperature-dependent heterophylly (Nakayama et al., 2014). However, despite the availability of transcriptomic data for several heterophyllous plants (van Veen et al., 2013; He et al., 2018; Han et al., 2021; Koga et al., 2021; Horiguchi et al., 2023), the molecular mechanisms underlying heterophylly are still largely unknown.

*Hygrophila difformis* (Acanthaceae), also known as water wisteria, is an amphibious plant that exhibits heterophylly by developing deeply lobed, complex leaves under submergence and simple leaves in terrestrial conditions (Fig. 1A). *H. difformis* has recently gained the attention of researchers as a promising new model for heterophylly studies, due to its beneficial traits such as easy gene transformation, fast propagation, and conspicuous leaf shape change in response to various environmental stimuli (Li et al., 2017; Li et al., 2020). It also allows us to explicitly address the interaction between environmental responses, morphological features, and variation in gene expression in an ecological context (Li et al., 2021). Previously, we showed that *SHOOT MERISTEMLESS* (*STM*) participates in the heterophylly regulation of *H. difformis* (Li et al., 2022), but key genes and molecular mechanisms underlying this phenomenon are still unclear. The genus *Hygrophila* comprises nearly 90 species, many of which are amphibious and display phenotypic plasticity (Fig. S1).

**Figure 1.**
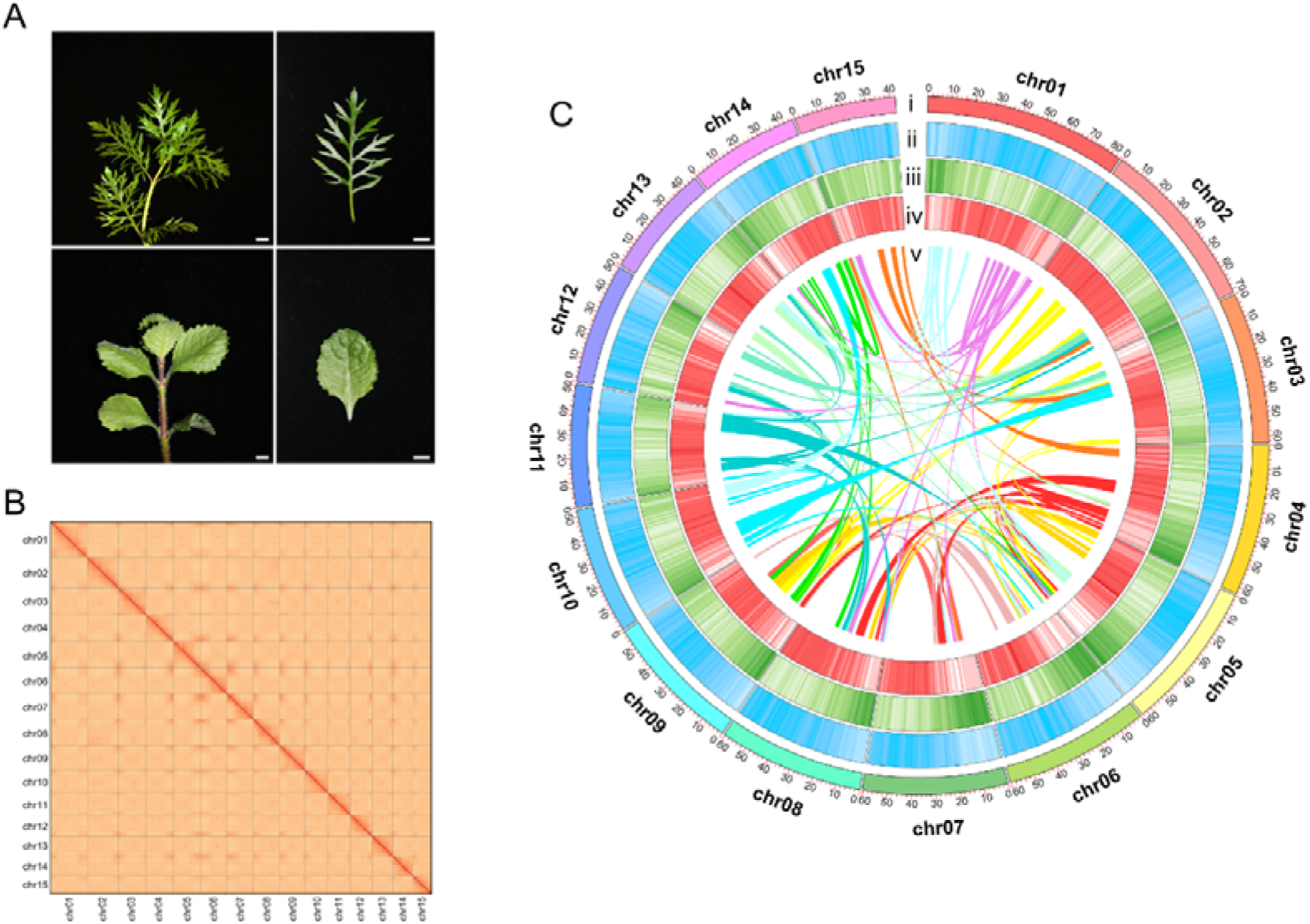
Genome assembly of *Hygrophila difformis*. (A) Morphology of the submerged plant (top, left), submerged leaf (top, right), terrestrial plant (bottom, left), and terrestrial leaf (bottom, right) of *H. difformis*. Bars = 1□cm. (B) High-throughput chromosome conformation capture map of the *H. difformis* genome showing a strong intra-chromosomal interaction signal. (C) Overview of *H. difformis* genome assembly. Chromosome (i), repeats (ii), gene density (iii), GC ratio (iv), and collinearity blocks (v) are displayed. The colour density from light to dark indicates the density of indicated features from low to high.

*H. difformis*, however, displays the most conspicuous form of heterophylly. *Hygrophila* includes species that are grown as ornamental plants (Karatas et al., 2013), used as medicinal herbs (Ingale et al., 2013), and even serve as models for photosynthetic studies (Botts et al., 1990; Horiguchi et al., 2021). However, at present, there are no genomic resources available for this genus.

Here, we report a chromosome-scale genome of *H. difformis*, assembled by integrating PacBio, Illumina, and High-throughput Chromosome Conformation Capture (Hi-C) sequencing technologies. We performed genome and transcriptome analyses to reveal the molecular mechanisms underlying heterophylly in response to the amphibious environment. We also identified and validated *LATE MERISTEM IDENTITY 1* (*LMI1*) as a candidate gene for leaf shape regulation. Our study provides the first genome and transcriptome for the genus *Hygrophila*, establishing *H. difformis* as a novel ecological model for investigating environmental impacts on plant phenotypes. This study offers a comprehensive framework and functional data for future genetic, genomic, and molecular studies of plant phenotypic plasticity. We believe that our work will attract more attentions to the plant phenotypic plasticity and environmental adaptation, and use *H. difformis* as an emerging model for plant Eco/Evo/Devo studies.

## Results

### Genome sequencing, assembly, and annotation

We previously reported that *H. difformis* reproduces through vegetative ways (Li et al., 2017), may be infertile and likely a triploid. Here we generated, 142.27 Gb sequencing data (genome coverage: 150 ×), with 52,199,178,926 *k*-mers and a *k*-mer depth of 54 (supplementary table S1, Fig. S2), which estimated a genome size of 913.38 Mb. The *K*-mer analysis confirmed that *H. difformis* is triploid (Fig. S2), as its two peaks have triple values (*K*-mer depth =18 and 54) (Ranallo-Benavidez et al., 2020). We also performed karyotype analysis and cytogenetic studies, which showed the presence of 45 chromosomes (2n = 3X = 45) (Fig. S3). We further evaluated the genome size by flow cytometry, and the detected genome size was 0.89 Gb (Fig. S4), slightly smaller than the *k*-mer estimated genome size (supplementary table S1). The genome of *H. difformis* was sequenced, separated, and assembled to the haploid genome using the PacBio Sequel platform (Zhou et al., 2021). The initial genome assembly consisted of 2,752 contigs covering 914.25 Mb. Using the Hi-C seq platform (Burton et al., 2013; Durand et al., 2016), we obtained 15 pseudochromosome sequences covering 871.92 Mb (95.37%), with a Scaffold N50 of 60.43 Mb (supplementary table S2-4). BUSCO (Benchmarking Universal Single-Copy Orthologs) (Simao et al., 2015) analysis suggested that 95.50% of the total genes were recovered (supplementary table S5). Based on the Hi-C contig contact matrix heatmap and genome features (Fig. 1B, C), we verified that the clustering, ordering, and orientation of the contigs were successful.

We identified 36,099 protein-coding gene models (supplementary table S6) according to homology-based and *de novo* predictions and 90.62% (32,714) of them could be mapped with functional annotations using public databases (supplementary table S7). We also performed BUSCO analysis using the predicted protein sequences to assess the completeness of our annotation, which indicates that 94.40% of the total genes were recovered (supplementary table S5). These results suggest a successful assembly and annotation of major genic regions in the *H. difformis* genome.

Repetitive DNA sequences are a major component of eukaryotic genomes, reflecting their organization and evolution (Mehrotra and Goyal, 2014). Using a combination of homology-based searches and *de novo* prediction, we identified repetitive sequences accounting for 74.92% of the genome (supplementary table S8), which is higher compared to other reported related terrestrial species within Acanthaceae family members, *Andrographis paniculata* (53.3%) (Sun et al., 2019), *Strobilanthes cusia* (74.4%) (Xu et al., 2020), *Avicennia marina* (48.8%) (Ma et al., 2022), and the reported related aquatic species *Utricularia gibba* (42.18%) (Lan et al., 2017) within the order Lamiales. Among the repeat sequences in *H. difformis* genome, long terminal repeats (LTRs) were dominant (62.39%) (supplementary table S8), which is also higher compared to *U. gibba* (12.44%), *A. marina* (24.92%), *A. paniculata* (38.54%), and *S. cusia* (51.32%). These results suggest that the genome of *H. difformis* is relatively more repetitive than the genomes of other reported related species.

### Comparative genomics and genome evolution

To investigate the evolution of *H. difformis*, we compared its genome to fourteen representative species of diverse evolutionary status, including one monocot, *Oryza sativa* (Poaceae), and thirteen eudicots. We identified a total of 14,656 ortholog groups, of which 1,362 were unique to *H. difformis*. (Supplementary table S9). We identified 210 single-copy orthologous gene sets among the 15 genomes and subsequently constructed a phylogenetic tree based on the molecular clock (Kumar et al., 2017). The phylogenetic placement of these species is consistent with references therein (Kordyum and Mosyakin, 2020). Among the genomes of other Lamiales, we estimated the divergence times of around 69.8 Mya (Million years ago) between *O. europaea* and the other six species, 57.8 Mya between *B. hygrometrica* and the remaining five species, and 30.6 Mya between *A. paniculata* and *H. difformis* (Fig. 2A). The differentiation time of *H. difformis* was approximately 25.4–37.9 Mya, coinciding with the evolution and diversification of East Asian Flora (EAF) during the Oligocene (33.9 to 23.0 Mya) (Ma et al., 2019; O’Brien et al., 2020).

**Figure 2.**
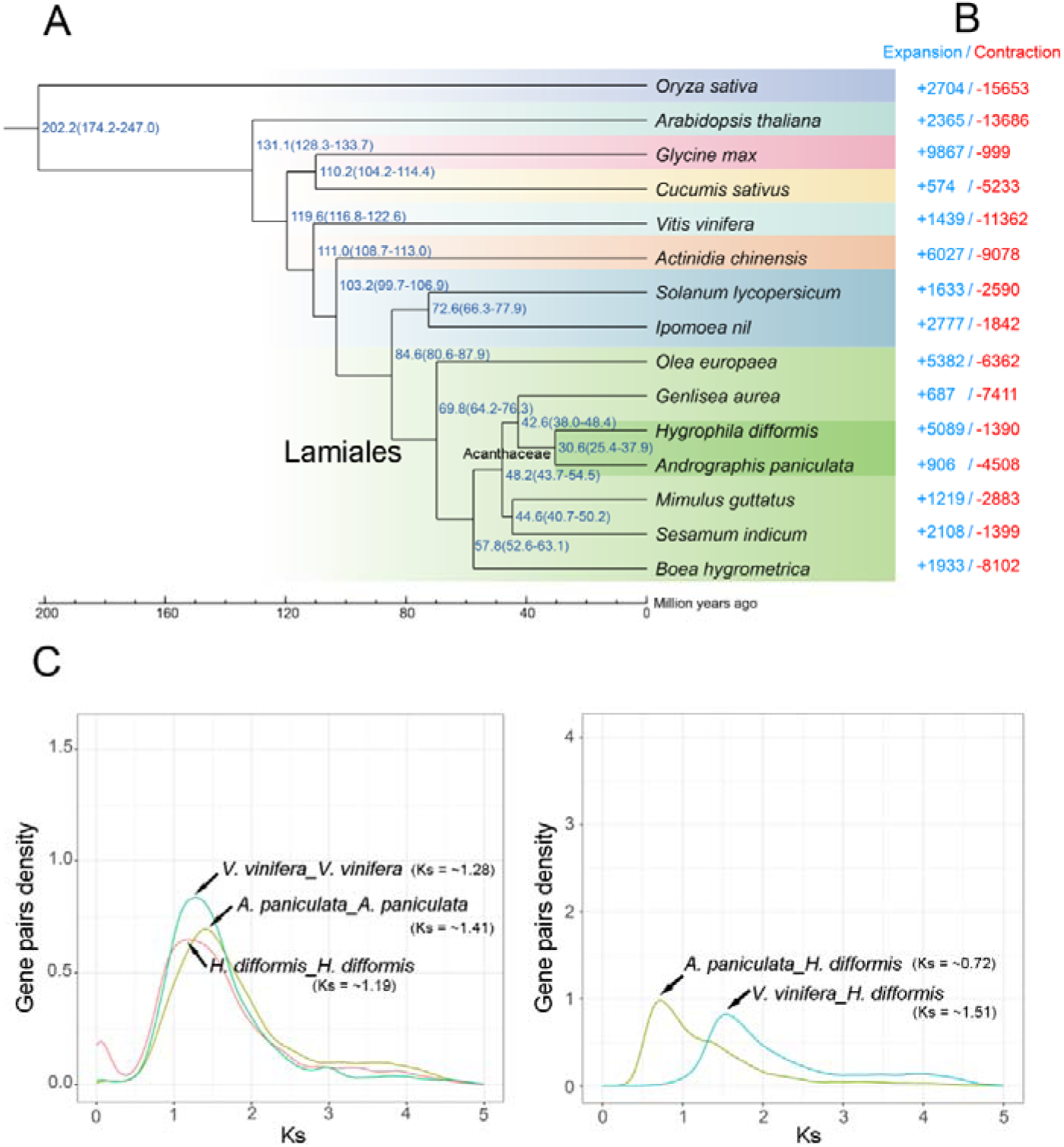
Evolutionary analysis of the *H. difformis* genome. (A) Phylogenetic relationships and times of divergence between *H. difformis* and other species. Different color backgrounds indicate species of diverse evolutionary status. Shallow green, Lamiales; Dark green, Acanthaceae. (B) Expansion and contraction of gene families. Red, contraction; Blue, expansion. (C) Distribution of average synonymous substitution levels (*Ks*) between syntenic blocks. These paralogs and orthologous gene pairs are among *H. difformis*, *A. paniculata* and *Vitis vinfera*. Left, paralogs; right, orthologs.

We then examined gene expansion and contraction in the phylogenetic analysis. We found that *H. difformis* had more gene expansion than its related species (Fig. 2B). A total of 5,089 ortholog groups were expanded in the *H. difformis* lineage, while 1,390 ortholog groups were contracted. GO pathway enrichment analysis showed that the significantly expanded (Q value < 0.01) genes were enriched in traits related to environmental adaptation, such as “tropism” (GO:0009606), “response to freezing” (GO: 0050826), and “response to blue light” (GO:0009637), and also in energy metabolisms and physiological processes, such as “carbohydrate metabolic process” (GO:0005975), “oxidation-reduction process” (GO:0055114) and “regulation of photosynthesis” (GO:0010109). We also detected expanded genes in the “hormone metabolic process” (GO:0042445) and “regulation of hormone levels” (GO:0010817), which may regulate the hormone levels and homeostasis in *H. difformis* (supplementary table S10). The significantly contracted (Q value < 0.01) genes were enriched in “carbon-oxygen lyase activity” (GO:0016838), “terpene synthase activity” (GO:0010333), and some other pathways (supplementary table S11).

Whole genome duplication (WGD) events are major evolutionary forces and have been associated with macroevolutionary changes and evolutionary innovations that happened relatively frequently in plants (Ren et al., 2022). Many genera of Lamiales underwent additional WGD events after the gamma event (Godden et al., 2019). To study the evolution of *H. difformis* within Acanthaceae, we characterized the synonymous substitution divergence (*Ks*) of each collinear gene pair within a genome or between genomes for WGD analysis between *H. difformis* and other reported species (Fig. 2C, Fig. S5). The *Ks* distribution of paralogs showed that *H. difformis*, grape (*Vitis vinifera*, the outgroup of Acanthaceae) and *A. paniculata* have only one peak at *Ks* = ∼1.19, ∼1.28 and ∼1.41, respectively (Fig. 2C). It should be mentioned that *H. difformis* likely experienced a recent WGD event that led to a triploid karyotype, which was not captured in this analysis. The orthologous distribution peaks for *H. difformis*–*A. paniculata*, *H. difformis*–*V. vinifera* also correspond to *Ks* values of ∼0.72 and ∼1.53, respectively (Fig. 2C). These results showed that the *H. difformis* genome may only experience the ancient whole-genome triplication (γ), which is similar to its close-related species *A. paniculata* and *S. cusia* within Acanthaceae (Sun et al., 2019; Xu et al., 2020). We revealed that the *Ks* distribution peak within *V. vinifera* was less than the *Ks* peak of *H. difformis*-*V. vinifera*, which indicated that the γ event occurred after their differentiation (Fig. 2C). We also analyzed the *Ks* distribution of recent reported Acanthaceae species *A. marina* and *S. cusia* (Fig. S5). In contrast, *Ks* distribution peaks of *H. difformis*–*S. cusia* and *H. difformis*–*A. paniculata* were less than the *Ks* peak within *H. difformis*, *S. cusia* and *A. paniculata* (Fig. 2C, Fig. S5), which indicated that these species underwent a common γ event. In addition, *A. marina* (2n= 64) experienced a special recent WGD (*Ks* = ∼0.49) (Fig. S5), which is similar to the recently published research (Ma et al., 2022).

### Morphological, physiological, and molecular adaptation of H. difformis to the terrestrial or submerged environment

Heterophyllous plants grown in submerged conditions often develop filiform, dissected or linear leaves that lack cuticle and stomata, while the terrestrial leaves are thicker, with differentiated mesophylls, and are cutinized with stomata (Koga et al., 2021; Li et al., 2021). We found that leaf area, dissection index (DI), and epidermal cell complexity of *H. difformis* were significantly increased in submerged conditions (Student’s *t*-test, *P* < 0.01), while stomatal density, vein density, and chlorophyll content were significantly increased in terrestrial conditions (Student’s *t*-test, *P* < 0.01) (Fig. 3A, S6), which is similar to other heterophyllous plants (Kim et al., 2018; Koga et al., 2021).

**Figure 3.**
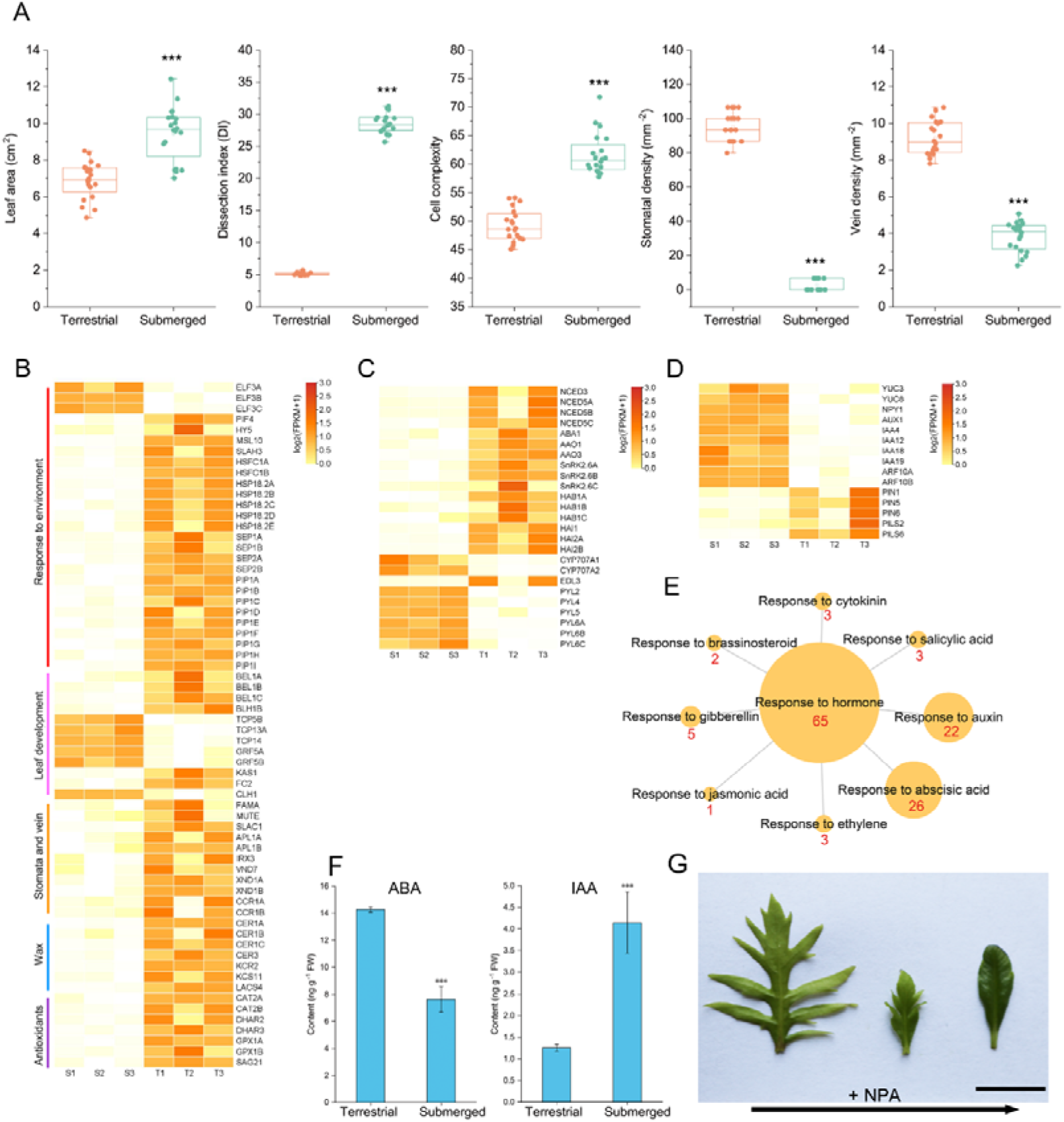
Phenotype and transcriptomic analyses of *H. difformis* under terrestrial or submerged conditions. (A) Morphological parameters (leaf area, dissection index, cell complexity, stomatal density, vein density, and chlorophyll content) of *H. difformis* in terrestrial or submerged conditions. Data from twenty biological replicates for each condition. ***, significant differences detected by Student’s *t*-test (*P* < 0.01). (B) DEGs (false discovery rate [FDR] < 0.05 and expression-level log_2_ [fold-change] > 1) involved in environmental response, leaf development, stomatal density, vein density, wax and antioxidants. Relative gene expressions were scaled to see their differences. (C) DEGs that are involved in ABA metabolism and signalling. Relative gene expressions were scaled to see their differences. (D) DEGs that are involved in auxin metabolism and signalling. Genes’ relative expressions were scaled to see their differences. (E) Diagram for phytohormone-related categories showing differential expression in terrestrial or submerged conditions. The number of genes is in red and is represented by the relative size of the circle corresponding to each gene category. (F) ABA and IAA content in shoots of *H. difformis* under terrestrial or submerged conditions. Error bars represent ± SD. Asterisks indicate significant differences (Student’s *t*-test: ***, *P* < 0.001). Data presented here are the means of three independent experiments. (G) Effect of NPA on *H. difformis* heterophylly. Photos show the emerging leaves after 10 μM NPA treatment for 1 month. The oldest is on the left, and the youngest leaf is on the right. The photo is of one representative from ten independent plants treated with NPA. Bar = 1LJcm.

To elucidate the molecular mechanisms underlying the phenotypic plasticity in *H. difformis*, we performed RNA-seq analysis on both submerged and terrestrial shoots using annotations based on our genomic data. A total of 6,226 differentially expressed genes (DEGs) (false discovery rate [FDR] < 0.05 and expression-level log_2_ [fold-change] > 1) were identified, with 4,100 upregulated genes and 2,126 downregulated genes in terrestrial shoots. We then analyzed the DEGs that are involved in environmental sensing or response (Fig. 3B). The thermosensor gene *EARLY FLOWERING 3* (*ELF3*) (Jung et al., 2020), a negative regulator of thermomorphogenesis, is downregulated in terrestrial conditions (Fig. 3B). *ELF3* has been shown to function antagonistically with *PHYTOCHROME-INTERACTING FACTOR4* (*PIF4*) in thermomorphogenesis (Raschke et al., 2015), and we also observed an inverse expression pattern of *PIF4* in *H. difformis* grown aerially. The proton sensor gene *SLAC1 HOMOLOGUE 3* (*SLAH3*) (Lehmann et al., 2021), the mechanical stimuli sensor gene *MSCS-LIKE 10* (*MSL10*) (Tran et al., 2021), and the *PLASMA MEMBRANE INTRINSIC PROTEIN* (*PIP*) aquaporin genes (Yaneff et al., 2015) were also upregulated in terrestrial conditions.

Morphological changes in leaf shape are the most obvious characteristics of heterophyllous plants. The antagonistic interaction of *KNOX1* and *BELL1-LIKE HOMEOBOX* (*BLH*) genes modulate leaf development, with *KNOX1* overexpression enhancing leaf lobing (Hareven et al., 1996) and the *BLH* overexpression reducing leaf serration (Kumar et al., 2007; Kimura et al., 2008). We observed increased expression of *BLH* genes in terrestrial shoots, which may underlie the leaf simplification observed in terrestrial conditions. *TfTCP13* (from *Torenia fournieri*) overexpression in *Arabidopsis thaliana* leads to narrower leaves (Zhang et al., 2021), and we detected increased expression of *TCP13* in submerged shoots of *H. difformis*. *TCP14* expression was reduced in the terrestrial leaves of *H. difformis*, mimicking the broad, short leaf phenotype induced by the dominant negative construct *TCP14::SRDX* in *A. thaliana* (Kieffer et al., 2011)*. GROWTH-REGULATING FACTORs* (*GRFs*) control leaf size in *A. thaliana* (Debernardi et al., 2014) and *GRF5* was elevated in the submerged shoots of *H. difformis*, which may account for the increased leaf size in submerged conditions (Fig. 3A, B). Genes involved in chloroplast development and chlorophyll biosynthesis are also elevated in terrestrial shoots (Fig. 3B, S6).

*MUTE*, *FAMA*, and *SLOW ANION CHANNEL-ASSOCIATED 1* (*SLAC1*), which regulate stomatal development and movement, are upregulated in terrestrial shoots, possibly facilitating stomatal formation in terrestrial leaves. Although *EPIDERMAL PATTERNING FACTOR-LIKE9* (*EPFL9*/*STOMAGEN*) is a positive regulator of the stomatal development (Hunt et al., 2010), it is often absent in aquatic plants, including *Z*. *marina*, *S*. *polyrhiza*, *Nymphaea colorata*, and *Wolffia australiana* (Park et al., 2021). However, we identified its putative homologous gene (Hdi026079) in *H. difformis*. Interestingly, its expression was not detected in either terrestrial or submerged shoots, suggesting this gene might be a pseudogene or it acquired a different function in *H. difformis*.

Genes involved in vascular development, lignin biosynthesis, and wax biosynthesis are upregulated in terrestrial shoots (Fig. 3B), which may enhance terrestrial acclimation. We also observed increased expression of genes responsive to oxidative stress in the terrestrial environment, which is in agreement with our finding that reactive oxygen species (ROS) accumulate in terrestrial leaves of *H. difformis* (Fig. 3B, S7).

### Roles of auxin and ABA in terrestrial and submerged acclimation

The role of phytohormones in the environmental control of morphogenesis has been reviewed recently (Nakayama et al., 2017; Li et al., 2019). We detected a large number of DEGs involved in the “response to hormone”, especially related to the “response to auxin” and “response to abscisic acid” (Fig. 3E). Our previous study showed that exogenous ABA can induce leaf simplification in *H. difformis* (Li et al., 2017). Here, we observed increased expression of genes related to ABA biosynthesis (e.g., *NINE-CIS-EPOXYCAROTENOID DIOXYGENASE* [*NCED*], *ABA DEFICIENT1* [*ABA1*], and *ALDEHYDE OXIDASE* [*AAO*]) and ABA signaling-related genes in terrestrial shoots, consistent with increased ABA content observed in terrestrial shoots (Fig. 3C, F). *CYP707A*, which encodes an ABA-degradation enzyme, was reduced in terrestrial conditions, as were the *PYRABACTIN RESISTANCE 1-LIKE* (*PYL*) genes. The expression pattern of these genes was consistent with their orthologs of *C. palustris* (Koga et al., 2021), suggesting similar ABA pathways regulate heterophylly in different amphibious species.

We also observed increased expression of auxin biosynthesis gene *YUCCA* (*YUC*) and the *YUC*-related *NAKED PINS IN YUC MUTANTS 1* (*NPY1*) in submerged shoots, along with increased auxin contents (Fig. 3D, F). *INDOLE-3-ACETIC ACID* (*IAA*)-*INDUCIBLE* and *AUXIN RESPONSE FACTOR 10* (*ARF10*) were also elevated in submerged conditions. Interestingly, the auxin influx transporter *AUXIN RESISTANT 1* (*AUX1*) was upregulated in submerged conditions, while the auxin efflux carriers *PIN-FORMED* (*PIN*) and *PIN-LIKES* (*PILS*) were downregulated.

To further examine the role of auxin transport in *H. difformis*, we treated submerged plants with the polar auxin transport inhibitor naphthylphthalamic acid (NPA), which caused a gradual reduction in leaf dissection and eventually the formation of oval leaves with smooth margins (Fig. 3G). Previous studies have reported the roles of ABA, ethylene (Kuwabara et al., 2003), GA, and cytokinin (Nakayama et al., 2014) in heterophylly. Here, we demonstrate for the first time that auxin metabolism and transport are also implicated in the regulation of heterophylly in *H. difformis*.

### Expression patterns of DEGs under diverse ecological factors

A previous study revealed expression patterns of candidate genes in the heterophyllous plant *C. palustris* under different phytohormone treatments (Koga et al., 2021). However, the response of heterophyllous plants to more natural and diverse environmental signals remain unclear. To address this, *H. difformis*, plants were grown in four sets of ecological factors (submerged/terrestrial, normal/low light density, normal/low temperature, and high/normal humidity) and sampled for RNA-seq (Fig. 4A). We also measured morphological and physiological characteristics between these groups (Fig. S8), showing similarities and differences affected by diverse ecological factors. Our transcriptomic analysis identified a total of 6,226 DEGs in the submerged/terrestrial group (Fig. 3), 9,213 DEGs in the normal/low light density group, 2,851 DEGs in the normal/low-temperature group; and 4,065 DEGs in the high/normal humidity group (Fig. 4B, 4C). A total of 180 genes were identified as upregulated DEGs and 119 genes were identified as downregulated DEGs in all four comparisons (supplementary table S12-13). *NCED3*, *ETHYLENE-INSENSITIVE 3* (*EIN3*), *CYTOKININ OXIDASE 5* (*CKX5*), and *GA INSENSITIVE DWARF1A* (*GID1A*) genes, which are related to phytohormone pathways, were among those genes elevated in all groups (Fig. 4D). Genes related to antioxidants were also upregulated, while *NPY1* was downregulated in all groups (Fig. 4E). A previous study showed that expression and functional divergence of *SUGARS WILL EVENTUALLY BE EXPORTED TRANSPORTERS* (*SWEET*) genes contribute to the specific developmental strategy of *Euryale ferox* leaves (Wu et al., 2022). Here, we observed that three putative orthologs of *SWEET* were upregulated in these groups, implying their potential roles in heterophylly. We also identified nineteen upregulated TFs and eight downregulated TFs (Fig. 4F) from these common DEGs, which may regulate the acclimation of *H. difformis* to different ecological conditions.

**Figure 4.**
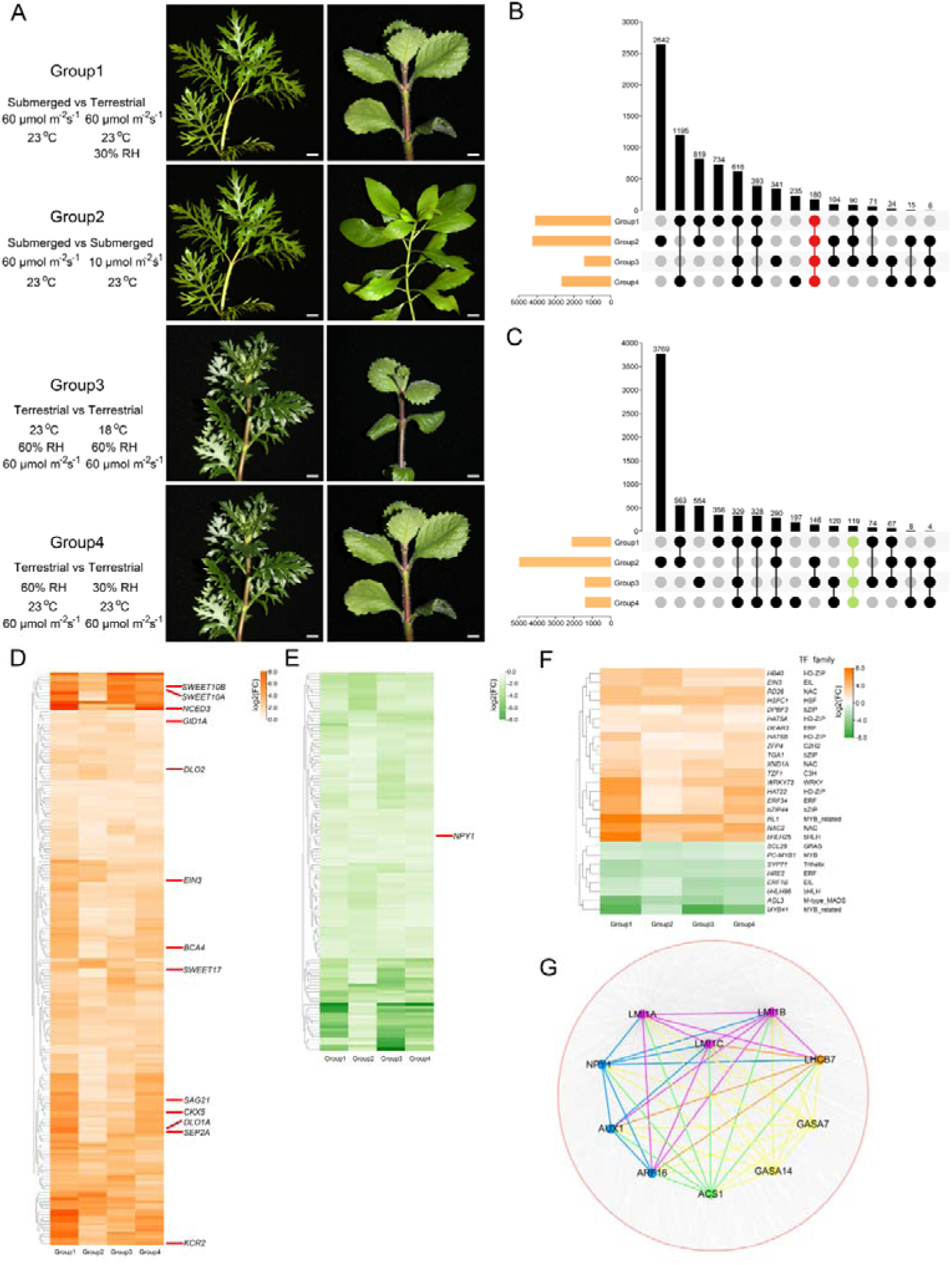
Phenotypic and transcriptomic analyses of *H. difformis* grown under diverse environmental treatments. (A) Phenotypes of *H. difformis* grown in different conditions. Group 1: submerged *vs* terrestrial plants. Group 2: Normal light density *vs* low light density. Group 3: Normal temperature *vs* low temperature. Group 4: high humidity *vs* normal humidity. Bars = 1cm. (B-C) Comparative transcriptome analyses of *H. difformis*. UpSet plots present the overlapping DEGs in 4 comparison groups. UpSet plot of DEGs upregulated compared to the control (B). UpSet plot of DEGs downregulated compared to the control (C). (D) Upregulated DEG expression profiles. (E) Downregulated DEG expression profiles. (F) Expression patterns of 27 putative TF genes. (G) Co-expression network reconstruction identified *LMI1* genes as being clustered with genes involved in auxin and GA signaling as well as ethylene synthesis and photosynthesis.

To further pinpoint key genes involved in heterophylly, co-expression analysis was conducted on a total of 15,295 DEGs (all DEGs that were changed in at least one group) and genes from 15 samples were filtered and clustered into ten modules using weighted gene co-expression network analysis (WGCNA) (Fig. S9). We focused on the yellow module which was associated with gene expression patterns and leaf complexity (Fig. S9, S10C). We identified three leaf development-related genes, putative orthologs of *LATE MERISTEM IDENTITY1* (*LMI1A*, *LMI1B*, and *LMI1C*), which were positively co-expressed with three auxin-related genes (*NPY1*, *AUX1*, and *ARF16*), two *GA-STIMULATED IN ARABIDOPSIS* (*GASA7* and *GASA14*) genes, the ethylene synthesis gene *ACC SYNTHASE1* (*ACS1*), and the photosynthesis-related gene *LIGHT-HARVESTING COMPLEX B7* (*LHCB7*) (Fig. 4G). It is known that the phenomenon of heterophylly involves morphological, anatomical and physiological adaptation (Horiguchi et al., 2019; Li et al., 2021). The candidate genes identified in our work regulating leaf formation, along with phytohormone pathways and physiological adaptation to environmental changes, might influence the heterophylly of *H. difformis*.

### HdLMI1 genes regulate the leaf shape of H. difformis in submerged conditions

*LMI1* and its putative orthologs have been studied in several terrestrial species and shown to have conserved roles in leaf shape diversity (Vlad et al., 2014; Andres et al., 2017). Here, we identified three putative paralogous *LMI1* orthologs in *H. difformis*, and examined their expression in different samples (Fig. 5A). These genes were highly expressed in humid and aquatic samples (A1-A3, and C1-C3), which had complex leaves (Fig. S10). In contrast, their expression was downregulated in simple-leafed groups (Fig. 5A).

**Figure 5.**
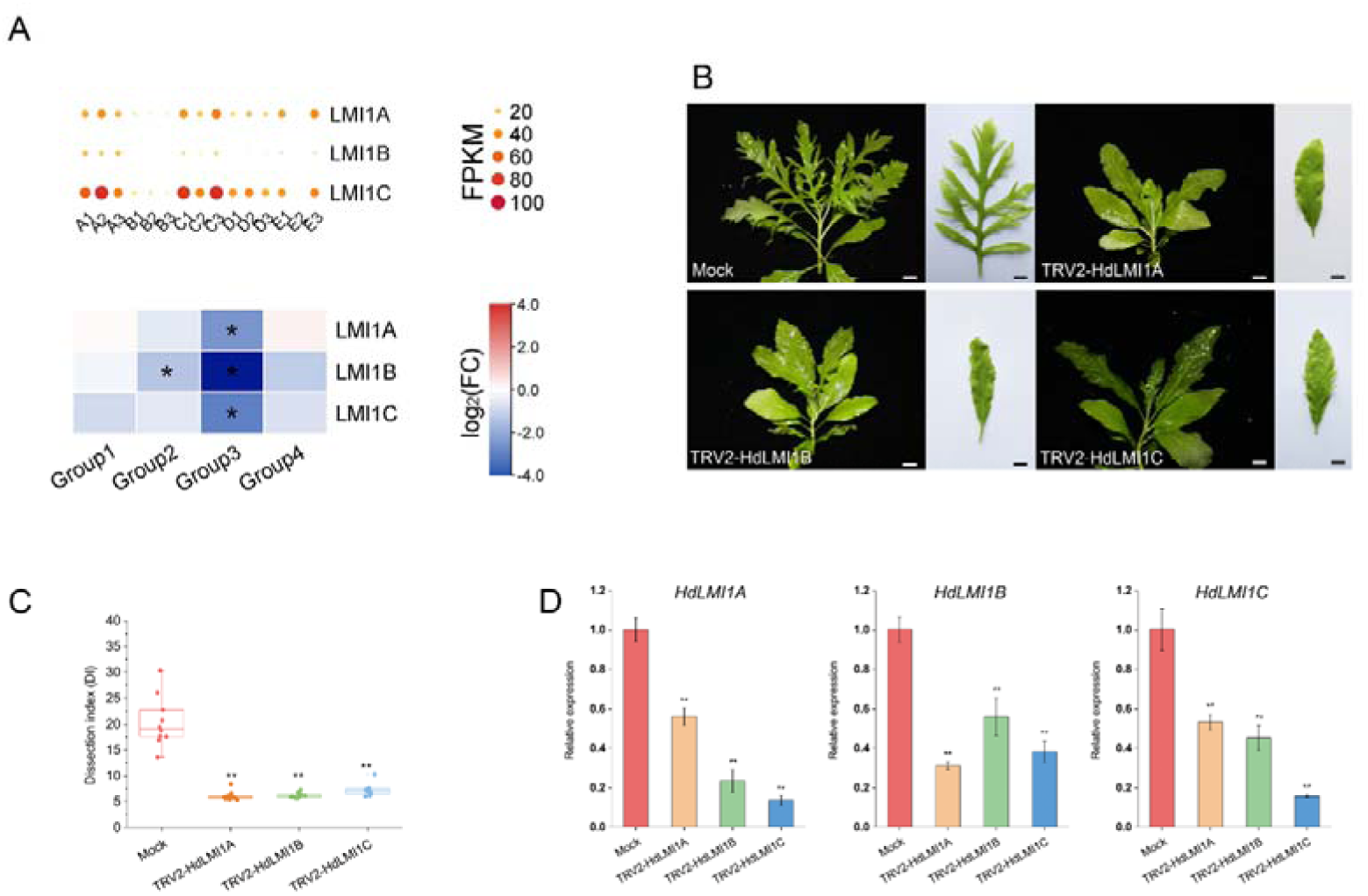
Expression of *LMI1* orthologs and function of *HdLMI1s*. (A) Relative expression of *LMI1* genes. Up, FPKM of *LMI1* genes in different environmental conditions. Down, Relative expression of *LMI1* genes in different groups. * Stars indicate DEGs (false discovery rate [FDR] < 0.05 and expression-level log_2_ [fold-change] > 1). (B) Phenotypes of whole plants and representative leaves in the mock, TRV2-HdLMI1A, TRV2-HdLMI1B, and TRV2-HdLMI1C transgenic plants grown in submerged conditions. Bars = 1 cm. (C) Dissection index in the mock, TRV2-HdLMI1A, TRV2-HdLMI1B, and TRV2-HdLMI1C transgenic plants grown in submerged conditions. Data from ten biological replicates for each, with significant differences between mock and transgenic plants detected by Dunnett’s test (**, *P* < 0.01). (D) Relative expression of *HdLMI1A*, *HdLMI1B*, and *HdLMI1C* in the mock, TRV2-HdLMI1A, TRV2-HdLMI1B, and TRV2-HdLMI1C transgenic plants. Error bars represent ± SD. Data presented here are the means from three independent experiments. Asterisks indicate significant differences (Dunnett’s test: **, *P* < 0.01).

To further examine the roles of *HdLMI1s* in regulating leaf development in *H. difformis*, we knocked down these genes using virus-induced gene silencing (VIGS) (Zhang et al., 2020) with the Tobacco rattle virus (TRV) constructs *TRV2-HdLMI1A*, *TRV2-HdLMI1B* and *TRV2-HdLMI1C*. Compared to mock plants, the *TRV2-HdLM*I1A, *TRV2-HdLMI1B*, and *TRV2-HdLMI1C* transgenic plants grown in submerged conditions were similar to each other, developing crinkled, complete, or shallowly serrated leaf margin (Fig. 5B). DI indicated that the mock plants had significantly higher leaf complexity than *TRV2-HdLMI1A*, *TRV2-HdLMI1B*, and *TRV2-HdLMI1C* transgenic plants (Dunnett’s test, *P* < 0.01) (Fig. 5C). In addition to their morphological features, these plants showed a significantly reduced expression of all three *HdLMI1s* compared to mock (Dunnett’s test, *P* < 0.01) (Fig. 5D). A previous study showed that orthologs of *LMI1* in *Medicago truncatula* cooperatively regulate leaf margin development (Wang et al., 2021). These results confirmed our hypothesis that *HdLMI1s* are important regulators of heterophylly in *H. difformis*.

## Discussion

Our knowledge of genetic regulatory networks that contribute to the variation of plant phenotypes has been advanced by the use of the latest technologies, but the mechanisms of phenotypic plasticity still remain elusive mostly due to the scarcity of suitable models (Hoffmann and Sgro, 2011; van Veen and Sasidharan, 2021). Our published genome and transcriptome of *H. difformis* provide the resources for further studies to expand the knowledge on adaptations that enable this plant to thrive in a highly fluctuating environment. The genome of *H. difformis* also offers a chance to investigate convergent evolution among heterophyllous species.

Many aquatic species became polyploid during speciation (Albert and Renner, 2020; Lu et al., 2021), and polyploidy was verified as a factor in plants expanding their ranges or invading new environments as polyploid progenies often possess novel traits (Cheng et al., 2021). Polyploidy and hybridization in aquatic or amphibious plants are also associated with complex evolutionary histories (Les and Philbrick, 1993; Prancl et al., 2014). For example, water caltrop (*Trapa natans*) (Lu et al., 2021) and *C. palustris* are tetraploid (Koga et al., 2021), while *Cabomba caroliniana* has polyploid (3×, 6×, 8×) and aneuploid variation between individuals (McCracken et al., 2013). The number of chromosomes in *Hygrophila* species is varied, such as in *H. spinosa* (2n = 32), *H. augustifolia* (2n = 44) and *H. balsamica* (2n = 34) (Govindarajan and Subramanian, 1983). We previously reported that *H. difformis* always reproduces through vegetative propagation (Li et al., 2017) and does not seem to produce viable seeds. Our genome survey and cytogenetic studies revealed 45 chromosomes, and further analyses suggest that the sequenced *H. difformis* is triploid (2n = 3x = 45). The family Lamiaceae exhibit widespread but asymmetrical gene duplication patterns and ancient polyploidy (Godden et al., 2019). For example, *Paulownia tomentosa* (Paulowniaceae) had 3 inferred WGDs, while there were 7-18 WGDs in Nepetoideae (Godden et al., 2019). Previous studies reported that *A. paniculata* (2n = 48) and *S. cusia* (2n = 32) only experienced one γ event (Sun et al., 2019; Xu et al., 2020). However, *A. marina* (2n = 64) was reported to have two WGDs, and the recent WGD was attributed to a novel specific adaptation to global warming that occurred during the palaeocene–Eocene maximum (Ma et al., 2022). Our analysis confirmed the recent WGD in *A. marina*, but did not detect the common γ event, which could be due to data or methodological differences. We also observed that *H. difformis* possessed the same γ event as *A. paniculata* and *S. cusia*, and lacked any additional WGD during its evolutionary history (Fig. 2E). Polyploidy is widely regarded as a common speciation strategy that has pronounced implications for plant evolution and ecology (Van De Peer et al., 2017). *H. difformis* appears to be a natural triploid. There are only four Acanthaceae species with genome information reported so far (Ma et al., 2022), and thus more data about species within the *Hygrophila* genus is essential for evolutionary studies. Future studies on comparing subgenomes by the haplotype-resolved genome assembly will also be helpful in this regard.

Our comparative genome data showed that the differentiation time of *H. difformis* is about 30.6 Mya (Fig. 2A), coinciding with the evolution and diversification of East Asian flora (EAF) during the Oligocene (33.9 to 23.0 Mya). It was recorded that the decline of atmospheric CO_2_ and global cooling culminated in the Early Oligocene Aridification event at 31 Mya (Wasiljeff et al., 2022), which is close to the differentiation time of *H. difformis*. We found significantly expanded (Q value < 0.01) genes of *H. difformis* were enriched in “response to freezing”, “regulation of photosynthesis”, and “carbohydrate metabolic process”, which may reflect the environmental response, physiological plasticity and metabolic adaptations to the changing environments (supplementary table S10). In addition, the formation of the Asian monsoon system in the Oligocene influenced the plant evolution and diversification (Ma et al., 2019). The seasonality of rainfall increased progressively, achieving modern monsoon-like wet summers and dry winters, by the early Oligocene (Chen and Su, 2020). Therefore, plant species must have adapted to the seasonality of rainfall and drought. For example, the habitat diversification (terrestrial and aquatic/marshland) within *Pogostemon* (Lamiaceae) is related to adaptations to seasonally changing climate that was triggered by the Asian monsoon in southern and southeast Asia (Yao et al., 2016). It is known that monsoon systems involved in seasonal rain and fluctuated water levels, and the water level change was regarded as the driver of heterophylly in many amphibious species (Wanke, 2011; Nakayama et al., 2014). *Hygrophila* species are widely spread in East Asia, South America, and Southern Africa (https://www.gbif.org/species/9795574), and many of which are amphibious plants (Fig. S1). However, due to lack of enough genome and fossil data of *Hygrophila*, it would be premature to conclude that the amphibious habitat and phenotypic plasticity of *H. difformis* are the innovation for climate change that happened in Oligocene.

The leaflet development in *C. hirsuta* depends on *REDUCED COMPLEXITY* (*RCO*), a homolog of *LMI1* that represses leaf flanks growth and promotes leaflets formation (Vlad et al., 2014). *RCO* originated from gene duplication in the Brassicaceae family and was lost in *A. thaliana*, leading to leaf simplification (Vlad et al., 2014). However, *LMI1* has diverse functions in different species. For instance, *LMI1* regulates the formation of simple serrated leaves in *A. thaliana*. The *lmi1* mutation resulted in decreased leaf serration but formed leaflets in *A. thaliana* (Saddic et al., 2006), leaf serrations in *M. truncatula* (Wang et al., 2021), and rapeseed (*Brassica napus*) (Hu et al., 2018), and leaf lobes in cotton (*Gossypium hirsutum*) (Andres et al., 2017). To assess the function of *LMI1* in heterophylly, we silenced *HdLMI1* genes by VIGS and observed reduced leaf complexity in *H. difformis* transgenic plants (Fig. 5). However, the three *HdLMI1* genes may have cross-suppressed together by the TRV constructs due to their sequence similarities, which points to functional redundancies among the *HdLMI1* genes. *In M. truncatula*, two *LMI1* paralogous are separately expressed in leaf margin tooth and sinus, and cooperatively regulate the leaf serrations (Wang et al., 2021). Thus, it is necessary to distinguish the effects of silencing each *HdLMI1* gene via CRISPR/Cas9-mediated genetic modifications. Moreover, heterophylly may involve epigenetic regulation such as non-coding RNAs, DNA methylation, nucleosome assembly (Nakayama et al., 2014; Li et al., 2021) etc., and more advanced analyses such as target mimicry, single-cell sequencing, and CRISPR/Cas9 genetic modification are warranted in future studies.

Auxin is a key phytohormone that orchestrates development and thus participates in plant evolution (Moon and Hake, 2011; Finet and Jaillais, 2012). Auxin transport is essential for the initiation of leaf primordia and the formation of leaf margins (Shani et al., 2006). In Arabidopsis, auxin transport disruption leads to the absence of serration in leaves, whereas *PIN1* localization determines the local auxin maxima at the tips of emerging serrations (Hay et al., 2006). In compound-leaved species auxin transport perturbation causes a dramatic simplification of leaf morphology (DeMason and Chawla, 2004; Wang et al., 2005). Although auxin accumulation has been detected in some waterlogged plants (Qi et al., 2019), its role in heterophylly-induction of amphibious plants remains elusive. Previous studies measured various phytohormones in *C. palustris* under submerged/terrestrial conditions (Koga et al., 2021), but they focused on the importance of ABA, GA and did not analyze the role of auxin in detail. Here, we show that auxin levels are significantly increased under submergence, and the genes involved in auxin metabolism, transport and response (e.g., *NPY1*, *AUX1*, *PIN1*) are differentially expressed (Fig. 3, 4). Moreover, we demonstrate that the auxin transport inhibition abolishes heterophylly in *H. difformis*. The interplay between of auxin maxima and *KNOX1* genes underlies the evolution of serrations in *A. thaliana* and leaflets in *C. hirsuta* (Canales et al., 2010).The *LMI1*-like and *KNOX1* genes also jointly control leaf development in both cotton and *C. hirsuta* (Chang et al., 2019; Wang et al., 2022). A recent study revealed that *LMI1* of *M. truncatula* regulates leaf margin development by activating auxin transporter genes (Wang et al., 2021). Thus, putative orthologs of *LMI1*, *KNOX1,* and the auxin transport genes may cooperate in *H. difformis* heterophylly regulation. In addition, *PIF*-mediated auxin metabolism regulates plant adaptation to various environmental factors (Sun et al., 2013; de Wit et al., 2014; Burko et al., 2022). In this study, we identified *PIFs* among the DEGs involved in heterophylly. Future research may establish *PIFs* as network hubs integrating various environmental factors and developmental signals in the heterophylly of *H. difformis*.

We identified numerous DEGs associated with environmental stimuli (Fig. 3B), which form complex networks with plant growth genes. For instance, ELF3 is a thermosensor that regulates rhythmicity and temperature responsiveness (Jung et al., 2020). Similarly, PIF4 directly activates the auxin biosynthetic gene *YUCCA8* (*YUC8*) to control hypocotyl elongation (Sun et al., 2012), while the petiole phenotypes in *elf3* mutants depend on PIF4-PIF5 mediated GA abundance (Filo et al., 2015). Heat shock proteins also modulate light and temperature signaling via the ELF3-PIF4 module to affect the hypocotyl growth (Zeng et al., 2023). *MSL10* participates in the wound-triggered early signal transduction pathway (Zou et al., 2016), while *SLAH3* respond to flooding stress in *A. thaliana* (Lehmann et al., 2021). In summary, genes related to environmental stimuli can regulate signal sensing and phytohormone metabolism, which may act as key links between external changes and internal responses in *H. difformis*.

Flooding stress (submergence or waterlogging) is a major challenge to many terrestrial plants, especially crops (Bailey-Serres et al., 2012) in the present era of rapid climate change. Plants have evolved two main strategies to cope with the lethal effects of submergence (Bailey-Serres and Voesenek, 2008; Akman et al., 2012; Muller et al., 2021). The “escape” strategy involves shoot elongation to reach the surface and restore air contact. In contrast, the “wait” strategy involves growth reduction and energy conservation to survive until the floods recede. Recent studies have mainly focused on the “escape” strategy in crops, especially in deep-water rice (Kuroha et al., 2018; Nagai et al., 2020). Interestingly, the aquatic plant *N. nucifera* also adopts the “escape” strategy during early submergence (Deng et al., 2022). Heterophyllous plants appears to have evolved a third strategy, “variation”, to adapt to constant changes in submergence and dehydration (Fig. 6). Understanding the mechanisms of heterophylly could provide novel avenues to engineer resilient crops and develop agricultural practices to mitigate the effects of climate change.

**Figure 6.**
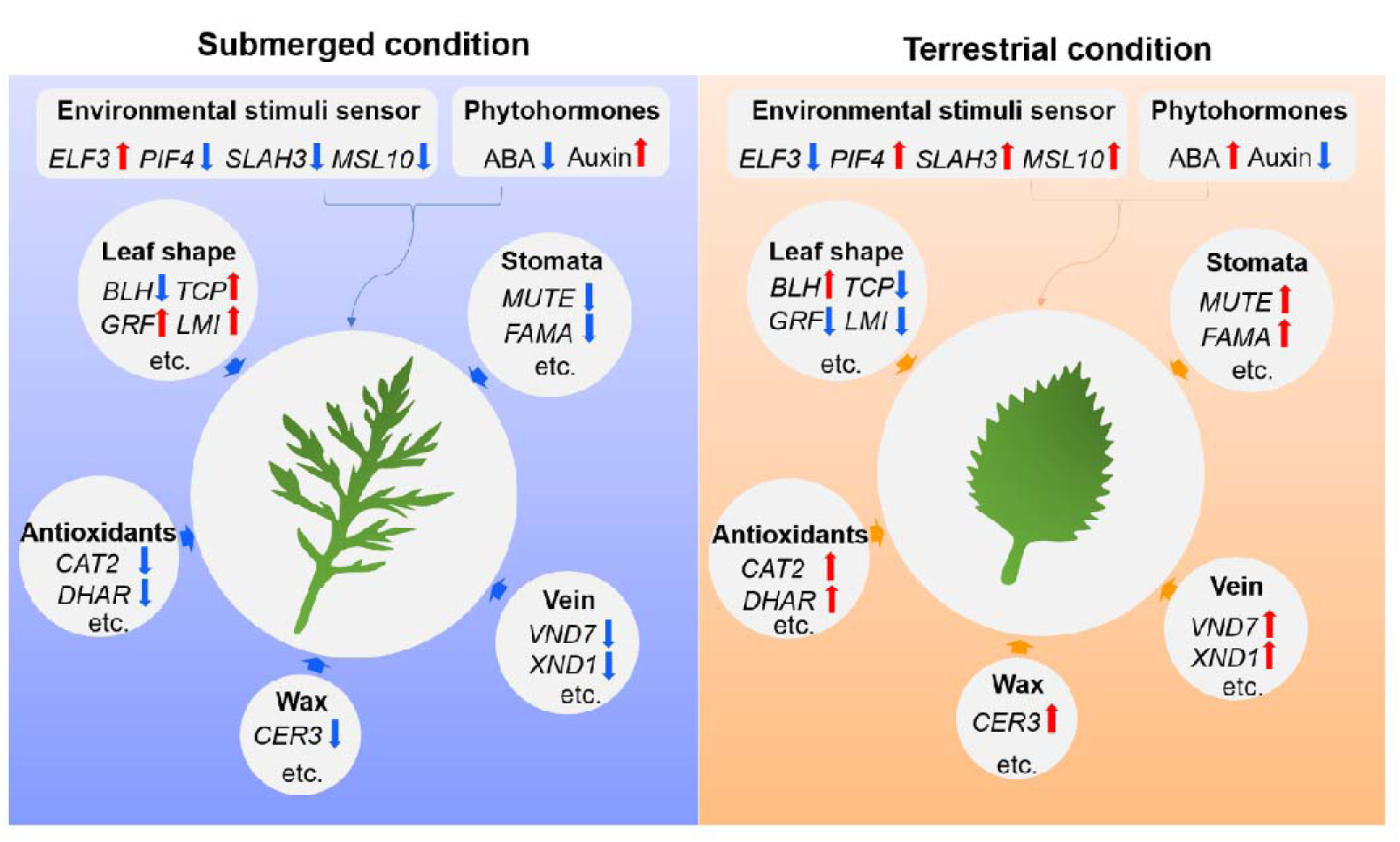
Schematic model of the regulation of heterophylly in *H. difformis*. The downregulated genes are indicated by blue arrows, whereas the upregulated genes are indicated by red arrows.

## Materials and Methods

### Plant materials and phenotypic analysis

The sequenced individual of *H. difformis* was collected from South China Botanical Garden, Chinese Academy of Sciences, Guangdong Province, China (N113.373781°, E23.18794°). For the analysis of submergence/emergence-treated samples, the shoots (at the same stage as our previous study (Li et al., 2022)) were harvested from plants grown under different experimental conditions (A: terrestrial plants grown at 23°C, 60% RH, 60 μmol m^-2^ s^-1^ illumination; B: terrestrial plants grown at 18°C, 60% RH, 60 μmol m^-2^ s^-1^ illumination; C: submerged plants grown at 23°C, 60 μmol m^-2^ s^-1^ illumination, D: submerged plants grown at 23°C, 10 μmol m^-2^ s^-1^ illumination, E: terrestrial plants grown at 23°C, 30% RH, 60 μmol m^-2^ s^-1^ illumination). For morphological analysis, mature leaves were photographed with a Canon EOS80D camera, and all light microscopy observations were performed under a Sunny EX20 light microscope and photographed with a ToupCam TP605100A digital camera. Images were integrated using MvImage media software (ToupCam). Leaf complexity was estimated based on the dissection index (DI), calculated as 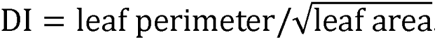, and epidermal cell complexity was also quantified a similar method of 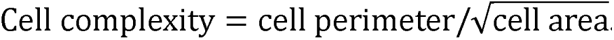. Stomatal and vein density are calculated as described (Li et al., 2017). All calculations were performed using ImageJ 1.47v (http://rsb.info.nih.gov/ij/). Statistical differences were determined using Student’s *t*-test or Tukey’s test, and data represent the results from at least twenty independent individuals.

### Estimation of genome size and ploidy

Short reads from the MGIseq2000 platform were filtered and used for genome size estimation by SOAPnuke (Chen et al., 2018). Initially, the adaptors were removed from the sequencing reads. Read pairs were then excluded if any single end has an average quality lower than 20, and subsequently, the ends of reads were trimmed if the average quality was lower than 20 in a sliding window size of 5 bp. Finally, read pairs with any end shorter than 75 bp were removed. These quality-filtered reads were used for genome size estimation. We generated the 17-mer occurrence distribution of sequencing reads from short libraries, and the genome size was estimated by 17-*K*-mer analysis based on the *K*-mer method (Manekar and Sathe, 2018) of Genome size = Kmer number/Kmer depth. The chromosome number of sequenced *H. difformis* was determined by a cytological method (Li et al., 2017). Root tips were fixed in an acetic-alcohol mixture (1:3) overnight. Cells were dissociated by 1 M hydrochloric acid at 60 °C for 1 h. After thorough washing with ultrapure water, they were stained by Modified Carbol fuchsin solution and squashed for chromosome number count. Genome size was estimated by flow cytometry as previously reported (Wang et al., 2014). We collected the young leaves from *H. difformis* and immediately carried out the flow cytometry (Dolezel and Bartos, 2005) to determine genome size by using *Sorghum bicolor* (BTX623, genome size = 0.71Gb) and *Zea mays* (Mo17, genome size = 2.18 Gb) as external standards.

### De novo sequencing and genome assembly

Tissue culture-raised plantlets from the *H. difformis* individual were utilized for the isolation of high molecular weight genomic DNA for genome sequencing. The quality and quantity of the extracted DNA were examined using a NanoDrop spectrophotometer (NanoDrop Technologies, Wilmington, DE, USA). Genomic DNA was isolated using the Qiagen DNeasy Plant Mini Kit (Qiagen, USA). We also collected a wide range of tissue samples (stem, shoot, root, flower, and leaves) from the same plant for RNA sequencing (RNA-seq) to support the genome assembly.

Two long-insert continuous long-read libraries (40 kb) were prepared following PacBio Sequel platform (Pacific Biosciences, Frasergen, Wuhan, China). The SMRT (Single Molecule Real-Time) library was constructed using the SMRTbell Express Template Prep kit 2.0 (Pacific Biosciences). Briefly, the DNA was carried into the first enzymatic reaction to remove single-stranded overhangs followed by treatment with repair enzymes. Subsequently, the ends of the double-stranded fragments were polished and tailed with T-overhang SMRTbell adapters and the SMRTbell library was purified using AMPure PB beads. The size distribution and concentration of the library were assessed using the FEMTO Pulse automated pulsed-field capillary electrophoresis instrument (Agilent Technologies, Wilmington, DE) and the Qubit 3.0 Fluorometer (Life Technologies, Carlsbad, CA, USA). Following library characterization, BluePippin system (Sage Science, Beverly, MA) was used to remove SMRTbells of ≤25 kb. After size selection, the library was purified with AMPure PB beads. Library size and quantity were assessed using the FEMTO Pulse and the Qubit dsDNA HS reagents Assay kit. Sequencing primer and Sequel II DNA Polymerase were annealed and bound, respectively, to the final SMRTbell library. SMRT sequencing was performed using a single SMRT Cell on the Sequel II System with Sequel II Sequencing Kit by Frasergen Bioinformatics Co., Ltd. (Wuhan, China). The draft assembly of the sequenced genome was assembled using MECAT2 (v20190226) with the default parameters (Xiao et al., 2017). The SMRT link toolkit was then performed to correct errors after the initial assembly of the genome.

To anchor hybrid scaffolds onto chromosomes, genomic DNA was also used to construct a Hi-C library. Briefly, samples were cross-linked under vacuum infiltration and the cross-linked samples were subsequently lysed. Endogenous nuclease was inactivated with 0.3% SDS, then chromatin DNA was digested by MboI (NEB), and marked with biotin-14-dCTP (Invitrogen) and then ligated by T4 DNA ligase (NEB). After reversing cross-links, the ligated DNA was extracted through QIAamp DNA Mini Kit (Qiagen) according to manufacturers’ instructions. Purified DNA was sheared to 300-to 500-bp fragments and were further blunt-end repaired, A-tailed and adaptor added, followed by purification through biotin-streptavidin–mediated pull-down and PCR amplification. Finally, the Hi-C libraries were quantified and sequenced on the Illumina Nova-seq platform (San Diego, CA, USA) or MGI-seq platform (BGI, China). We obtained sequencing data using a BGI MGISEQ-2000 platform on 150 PE mode. For anchored contigs, 381,400,686 clean read pairs were generated from the Hi-C library and mapped to the polished *H. difformis* genome using BWA (bwa-0.7.16) (Li and Durbin, 2010). Lachesis (Burton et al., 2013) was applied to order and orient the clustered contigs, and Juicer and Juicebox tool (v1.8.8) with default parameters (Durand et al., 2016) were used to correct the assembly error in the Hi-C assembled genome. Paired reads mapped to different contigs were used for the Hi-C-associated scaffolding. Self-ligated, non-ligated, and other invalid reads were filtered out. We applied 3D-DNA to order and orient the clustered contigs and performed Juicer to filter the sequences and cluster them. Juicebox with default parameters was applied to adjust chromosome construction manually. We also use purge haplotigs to remove heterozygosity. We finally anchored the scaffolds on fifteen chromosomes. In addition, the BUSCO pipeline was used to assess the completeness and accuracy of our assembled *H. difformis* genome as previous method (Simao et al., 2015).

### Prediction and annotation of protein-coding genes

We predicted the protein-coding genes of *H. difformis* using three methods: *ab initio*, homology-based, and transcriptomic prediction. AUGUSTUS (v3.3.1) (Stanke et al., 2006) and GlimmerHMM (v3.0.4) (Majoros et al., 2004) were used to perform *ab initio* gene prediction. We used EXONERATE (v2.2.0) (Slater and Birney, 2005) for the homology-based prediction. First, the protein sequences were aligned to the genome assembly and predicted coding genes using EXONERATE with the default parameters. For the transcriptomic prediction, we used RNA-seq analysis. We first assembled clean RNA-Seq reads into transcripts using trinity (Haas et al., 2013), and the gene structures were formed using PASA (Trapnell et al., 2010). Finally, MAKER (v3.00) (Cantarel et al., 2008) was used to integrate the prediction results. Gene functions were inferred according to the best match of alignments to the non-redundant database (NR) of the National Center for Biotechnology Information (NCBI) (https://www.ncbi.nlm.nih.gov/), TrEMBL (Boeckmann et al., 2003), SWISS-PROT (Boeckmann et al., 2003), and InterPro (Mitchell et al., 2015). Motifs and protein domains were annotated using PfamScan (Mistry et al., 2007) and InterProScan (http://www.ebi.ac.uk/InterProScan). We also performed BUSCO analysis (dataset: embryophyta_odb10 models) using the predicted protein to assess our annotation completeness of *H. difformis* genome.

### Annotation of repetitive sequences

The homology-based and *de novo* prediction methods are combined to identify the repeats in the genome. For homology-based analysis, we identified the known TEs within the *H. difformis* genome using RepeatMasker (open-4.0.9) (Tarailo-Graovac and Chen, 2009) with the Repbase TE library (Jurka et al., 2005). RepeatProteinMask searches were also conducted using the TE protein database as a query library. For *de novo* prediction, we constructed a *de novo* repeat library of the *H. difformis* genome using RepeatModeler (http://www.repeatmasker.org/RepeatModeler/), which can automatically execute two core *de novo* repeat-finding programs RECON (v1.08) (Bao and Eddy, 2002) and RepeatScout (v1.0.5) (Price et al., 2005) to comprehensively conduct, refine and classify consensus models of putative interspersed repeats for the *H. difformis* genome. Furthermore, we performed a *de nov*o search for long terminal repeat (LTR) retrotransposons against the *H. difformis* genome sequences using LTR_FINDER (v1.0.7) (Xu and Wang, 2007). Finally, we merged the library files of the two methods and used Repeatmaker to identify the repeat contents.

### Gene family identification and phylogenetic analysis

We selected 14 species to construct putative gene families: *Oryza sativa* (*Poaceae*), *Boea hygrometrica* (*Gesneriaceae*), *Mimulus guttatus* (*Phrymaceae*), *Olea europaea* (*Oleaceae*), *Sesamum indicum* (*Pedaliaceae*), *Andrographis paniculata* (*Acanthaceae*), *Genlisea aurea* (*Lentibulariaceae*), *Actinidia chinensis* (*Actinidiaceae*), *Arabidopsis thaliana* (*Brassicaceae*), *Cucumis sativus* (*Cucurbitaceae*), *Glycine max* (*Leguminosae*), *Ipomoea nil* (*Convolvulaceae*), *Vitis vinifera* (*Vitaceae*) and *Solanum lycopersicum* (*Solanaceae*), and their genome data were downloaded from the NCBI database (https://www.ncbi.nlm.nih.gov).

To cluster protein-coding genes into families, protein sequences from the longest transcript of each gene from *H. difformis* and other species were extracted and aligned to each other using the BLAST program with a maximal e-value of 1e^−5^. The genome sequences and corresponding protein sequences were downloaded from NCBI (https://www.ncbi.nlm.nih.gov/). Genes with identity < 30% or coverage < 50% were filtered out to exclude putative fragmented genes. The OrthoMCL (v14-137) (Li et al., 2003) method was used to cluster genes from these different species into gene families. To reveal phylogenetic relationships among *H. difformis* and other closely related species, protein sequences from 196 single-copy orthologous genes were used for phylogenetic tree reconstruction. The protein sequences of the single-copy orthologous genes were aligned with the MUSCLE (v3.8.31) program (Edgar, 2004), and the corresponding coding sequence alignments were generated and concatenated with the guidance of protein alignment. RAxML (v8.2.11) (Stamatakis, 2014) and TimeTree (Hedges et al., 2006) were used to construct the phylogenetic tree using the maximum likelihood method.

### Gene family expansion and contraction analysis

Based on the identified gene families and constructed phylogenetic tree with predicted divergence times of species, we used CAFÉ to analyze gene family expansion and contraction (Mendes et al., 2020). A random birth or death model in CAFÉ was used to study gene gain or loss in gene families across the phylogenetic tree, and an adjusted P value (Q value) was calculated for the genes of *H. difformis* to test the significant expansion or contraction (Fisher test, Q value < 0.01). These expansion and contraction genes were mapped to GO pathways for functional enrichment analysis, which was conducted using the enrichment methods. This method implemented hypergeometric test algorithms and the Q-value (FDR, False Discovery Rate) was calculated to adjust the p-value using the R package (https://github.com/StoreyLab/qvalue).

### Collinearity and WGD analysis

BLASTp (e-value <1e^-5^) was used to detect orthologous genes in *H. difformis*, *S. cusia* and *A. paniculata*, to assess the level of synteny between the species. Then, syntenic paralogous blocks were identified with MCScanX (Wang et al., 2012) integrated in TBtools software with default parameters, and visualized by TBtools (Chen et al., 2020). BLASTp (e-value <1e^-5^) was also used to detect orthologous pairwise, and MCScanX analysis was then performed for *Ks* evaluation of *V. vinifera*, *A. marina*, *H. difformis*, *S. cusia* and *A. paniculata*. Genome data and annotation of *V. vinifera* and *A. paniculata* were downloaded from the NCBI database (https://www.ncbi.nlm.nih.gov). Genome data and annotation of *S. cusia* were downloaded from its database (http://indigoid-plant.iflora.cn) (Xu et al., 2020). Genome data and annotation of *A. marina* were downloaded from its database (https://datadryad.org/stash/dataset/doi:10.5061/dryad.3j9kd51f5) (Friis et al., 2021). These analyses were performed using a haploid genome. We extracted all paralogous and orthologous gene pairs from syntenic blocks in those species to further estimate the *Ks* distribution and infer WGD dates by assuming a molecular clock (Schmutz et al., 2010).

### RNA sequencing, gene expression analysis, and co-expression analyses

Total RNA was extracted from shoots including leaf primordia of plants grown for a month under each condition were collected for RNA-seq, with each sample performing three biological replicates (Li et al., 2022). We used SOAPnuke (Chen et al., 2017) to filter the raw reads, and HISAT2 (Kim et al., 2015) to align filtered clean reads to the *H. difformis* genome. Read counts of these samples were calculated, and differential gene expression analysis was conducted using the R package DESeq2 (Love et al., 2014) (false discovery rate [FDR] < 0.05 and expression-level log_2_ [fold-change] > 1). For DEGs identification and naming, putative orthologs were identified by tBLASTn and BLASTp (e-value <1e-5) analysis to published Arabidopsis databases (https://www.arabidopsis.org/). To identify relationships between DEGs, co-expression networks were constructed using the WGCNA package (Langfelder and Horvath, 2008) integrated in TBtools software with default parameters, and DEGs were filtered according to their gene expression profiles (discarded genes that FPKM < 0.9). Cytoscape (Shannon et al., 2003) was used to visualize the network of genes related to phytohormones’ responses. Expression heatmaps were also generated using TBtools software (Chen et al., 2020).

### Quantification of phytohormones and NPA treatments

To measure phytohormone levels, plants were grown in growth chambers at 23°C under a white light flux density of 60 µmol m^−2^ s^−1^ for 1 month. Shoots including leaf primordia were dissected from the plants and frozen in liquid nitrogen immediately after sampling. Extraction and quantification of endogenous hormones (IAA and ABA) were performed as described previously (Nguyen et al., 2018). The experiments were conducted in triplicate from three independent plants. For naphthylphthalamic acid (NPA) treatments, plants grown in submerged conditions were shifted to an aquarium that contained a 10 µM NPA solution. After 1-month of treatment, emerging leaves were harvested for morphological analysis.

### ROS detection

Terrestrial and submerged mature leaves (at same stage, P6) were harvested and stained with DAB and NBT solution for H_2_O_2_ and O_2_^-^ analysis, respectively, as previously described (Chen et al., 2021). Leaves were harvested from twenty independent plants and immediately stained by H_2_O_2_ distribution (1% 3,3-diaminobenzidine, DAB), and O_2_^−^ distribution (0.1% nitro blue tetrazolium, NBT) overnight. Leaves were then rinsed in an ethanol series and recorded for analysis.

### Gene functional studies and expression analysis

To amplify the cDNA sequences of *HdLMI1A*, *HdLMI1B*, and *HdLMI1C*, total RNA was extracted from shoots including leaf primordia of plants grown for a month and then used to synthesize cDNA as described previously (Li et al., 2020). Amplified fragments of expected length were purified and cloned into a pEASY-T1 cloning vector (TransGen, China) for sequencing. The functions of *HdLMI1A*, *HdLMI1B*, and *HdLMI1C* were investigated using the VIGS method as previously described (Zhang et al., 2020). Specific fragments (171 bp for *HdLMI1A*, 268 bp for *HdLMI1B*, and 252 bp for *HdLMI1C*) were amplified, purified, and independently introduced into the pTRV2 vector (http://www.bt-lab.cn/) using a Trelief SoSoo Cloning Kit (Tsingke Biological Technology, China) to generate the constructs TRV2-HdLMI1A, TRV2-HdLMI1B, and TRV2-HdLMI1C. Recombinant vectors and empty vectors were then introduced into *Agrobacterium tumefaciens* strain LBA4404 by the liquid nitrogen freeze-thaw method. Empty vectors were introduced as mock controls. At least twenty wild-type individuals were used for each treatment. After two rounds of infiltration (7 and 14 days), plants were grown for 3 additional days in terrestrial conditions and then shifted to aquaria. After 20 days of growth in submerged conditions, representative leaves from ten individuals showing phenotypic changes were photo-documented for analysis. The silencing efficiency of VIGS treatments was determined by reverse transcription quantitative PCR (RT-qPCR) as previously described (Li et al., 2022). The cDNAs from individual developing leaves of the VIGS-treated plants were used as templates. The primers used in this study are provided in supplementary table S14. Significant differences in relative expression levels between mock and transgenic lines at *P* < 0.01 were evaluated using Dunnett’s test with the two-sided function in IBM SPSS version 24.

## Data availability

The genome dataset is available on GenBank of NCBI BioProject with the accession number PRJNA872356. Genome sequencing data (BGI seq reads, PacBio Sequel reads, and Hi-C interaction reads) and environmental treatment RNA-Seq reads are available in the NCBI Sequence Read Archive under accessions PRJNA855313 and PRJNA859677. Sequence data from this article can be found in the GenBank data library under the following accession number: *HdLMI1A* (OR296694), *HdLMI1B* (OR296695) and *HdLMI1C* (OR296696).

## Author contributions

G.J.L. and H.W.H. designed the experiments and wrote the manuscript; G.J.L., X.Y.Z., and J.J.Y. performed genome and transcriptome analyses; G.J.L. conducted environmental treatment experiments and collected the materials; S.Q.H., J.P., S.K., K.U.T., and I.H. improved the manuscript. All authors read and approved the final version of the manuscript.

## Funding information

This study was supported by the International Partnership Program of the Chinese Academy of Sciences (152342KYSB20200021), the National Natural Science Foundation of China (32101254), and the National Key Research and Development Program of China (2017YFE0128800). This work was also supported by the China Scholarship Council.

## Acknowledgements

We thank professor Shilin Chen of the Institute of Chinese Materia Medica, China Academy of Chinese Medical Sciences for kindly providing the genome annotation of *A. paniculata* and Dr. Markus Stetter, University of Cologne for critical reading of the manuscript. We thank Li Xie and Huan Wang of Wuhan Frasergen Bioinformatics Co., Ltd. for kindly providing the after service of our genome project.

## Notes

### Competing Interest Statement

The authors have declared no competing interest.

### Summary of Updates

new version

