## Supplemental Figures for "Water wisteria genome reveals environmental adaptation and heterophylly regulation in amphibious plants"

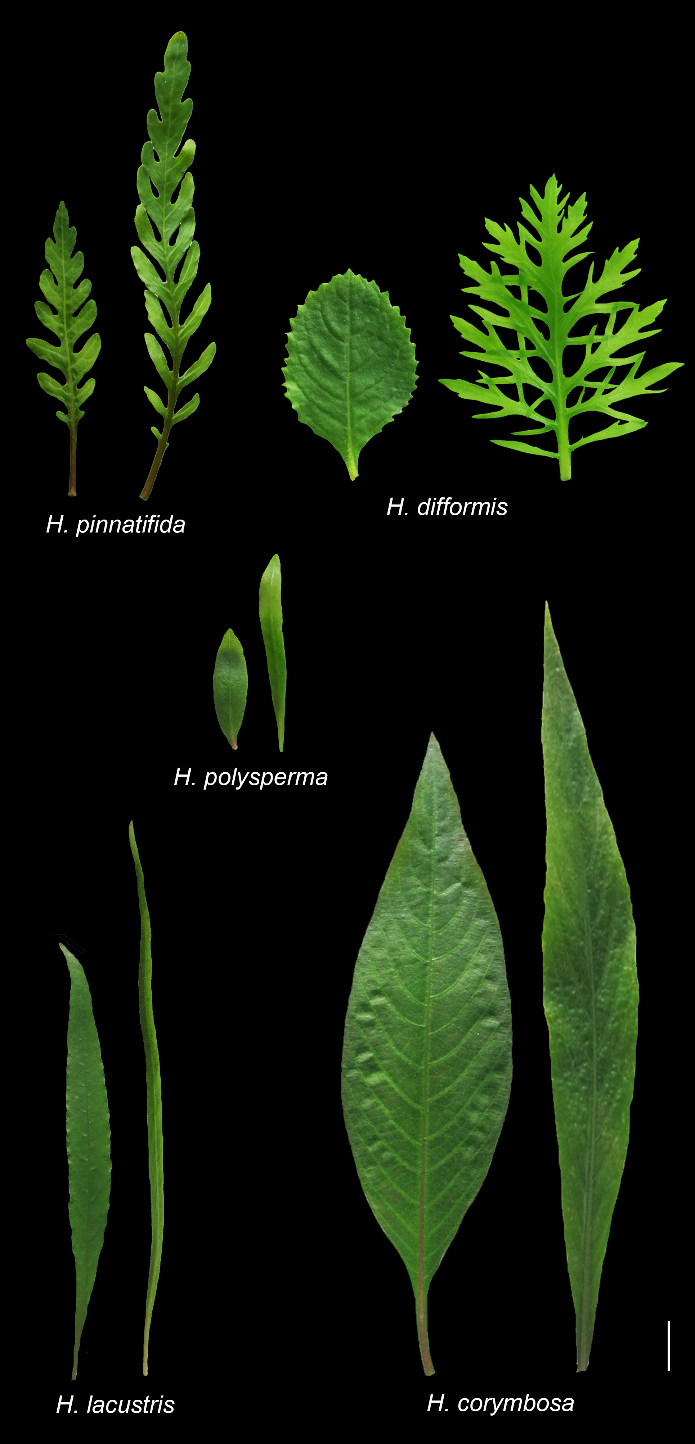

**Figure S1.** Phenotypic plasticity in the *Hygrophila* genus.

Photos show 5 pairs of leaves in *Hygrophila* species, which include *H. pinnatifida*, *H. difformis*, *H. polysperma*, *H. lacustris*, and *H. corymbose*. Left, a terrestrial leaf. Right, a submerged leaf. Leaf images were digitally extracted from the original pictures for comparison. Bar = 1 cm.

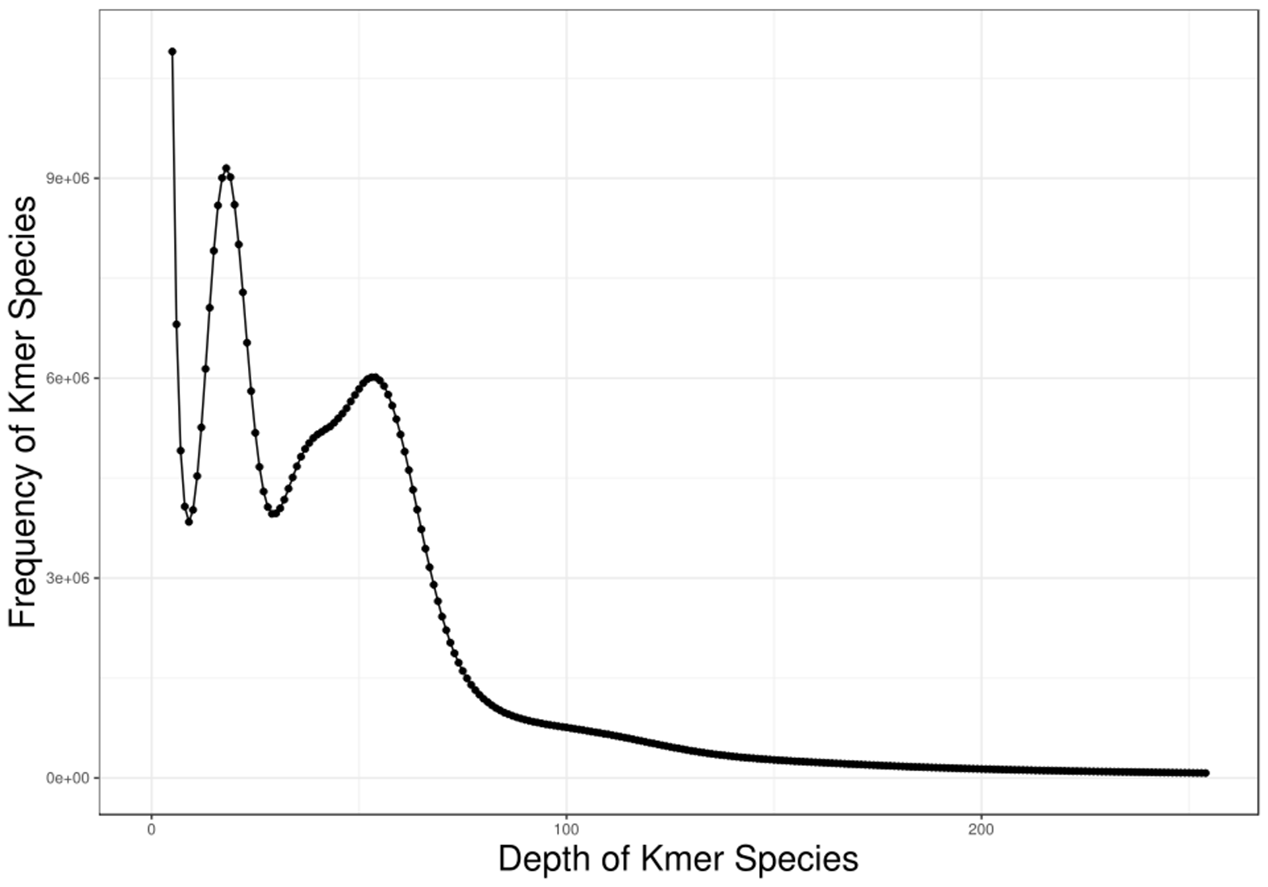

**Figure S2.** The 17-mer distribution of Illumina short-read data.

The figure shows the frequency of 17 *K*-mers which are 17 bp sequences from the reads (after filtering) of short-insert size libraries. The x-axis shows *K*-mer abundance. The y-axis shows the number of *K*-mers at a given abundance. Note that *H. difformis* has two clear peaks at *K*-mer depth =18 and 54.

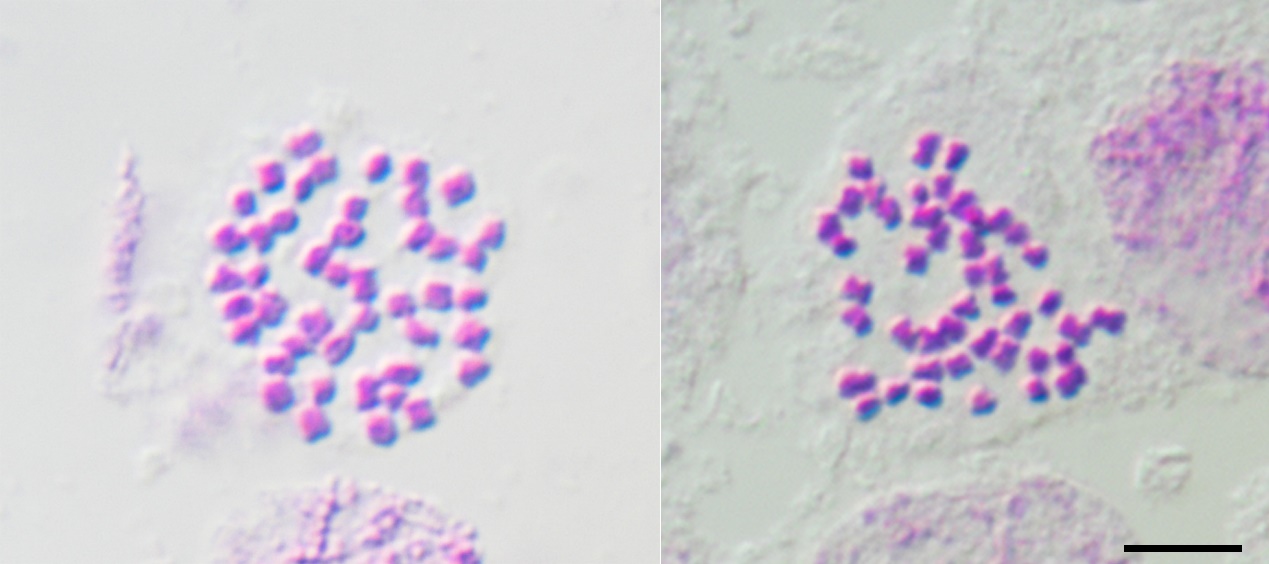

**Figure S3.** Mitotic metaphase chromosomes of the sequenced individual of *H. difformis*. Picture show two representatives of thirty individual cells. The chromosome number is 2n = 3X = 45. Bar = 10 μm.

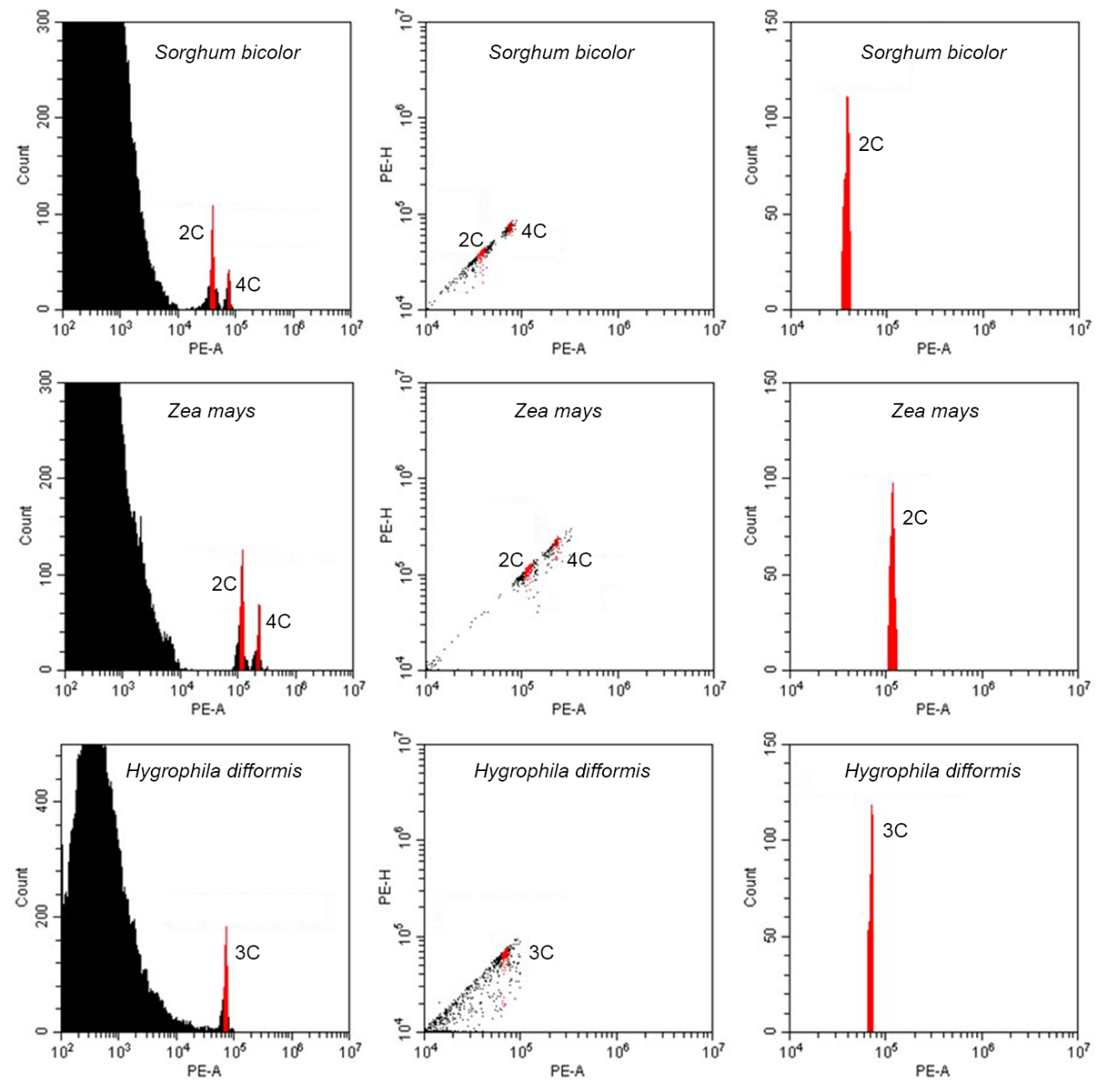

**Figure S4.** Flow cytometric analysis of the genome size of *H. difformis*.

*Sorghum bicolor* (genome size = 0.71 Gb) and *Zea mays* (genome size = 2.18 Gb) were used as external biological references. The genome size of *H. difformis* was estimated to be 0.89 Gb.

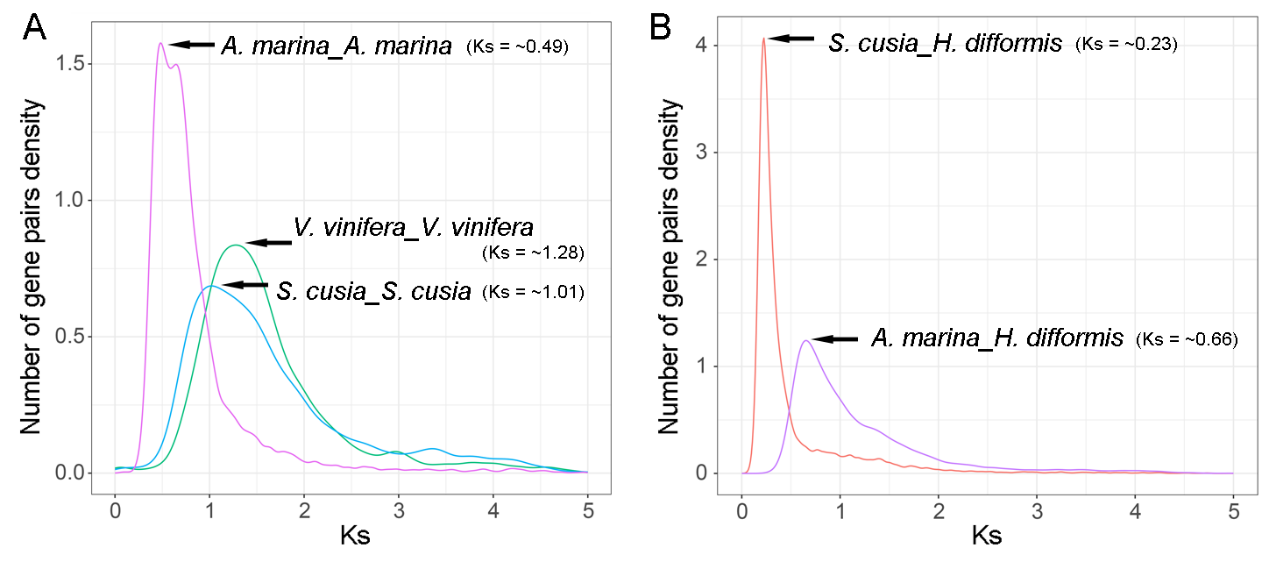

**Figure S5.** Distribution of average synonymous substitution levels (*Ks*) between syntenic blocks. These paralogs and orthologous gene pairs are among *Hygrophila difformis*, *Avicennia. marina*, *Strobilanthes cusia* and *Vitis vinfera*. Left, paralogs; right, orthologs.

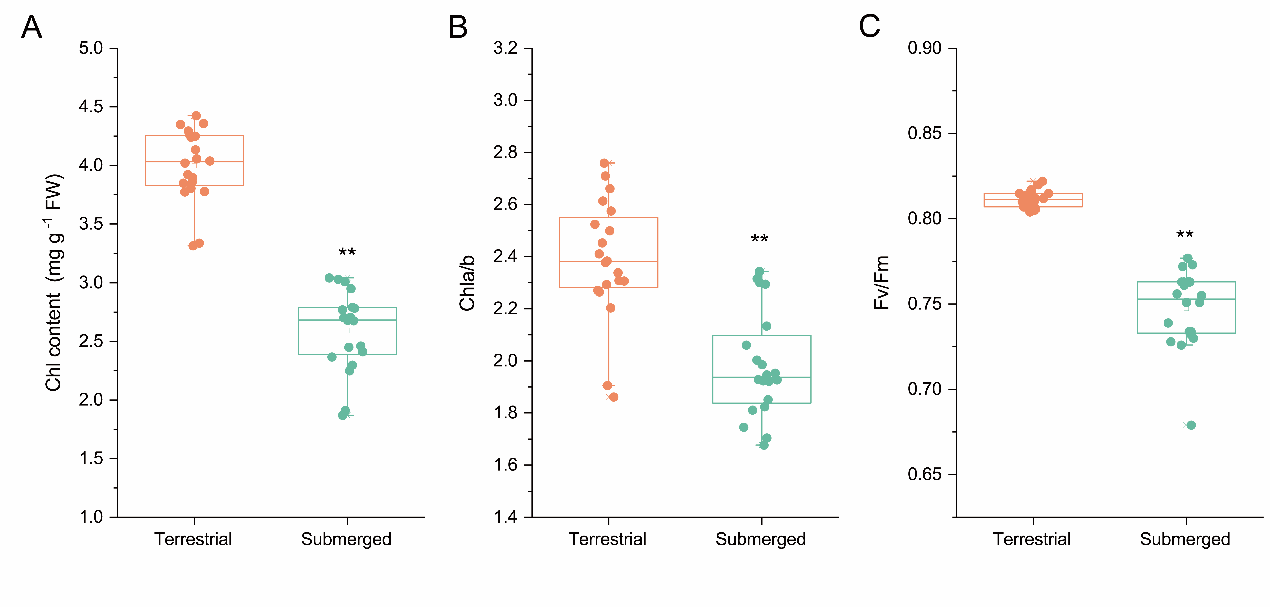

**Figure S6.** Photosynthetic physiology analysis of *H. difformis* in terrestrial and submerged conditions. (A) Chlorophyll content, (B) Chlorophyll a/b, (C) Fv/Fm. Data are from twenty biological replicates for each condition. **, significant differences detected by Student’s *t*-test (*P* < 0.01).

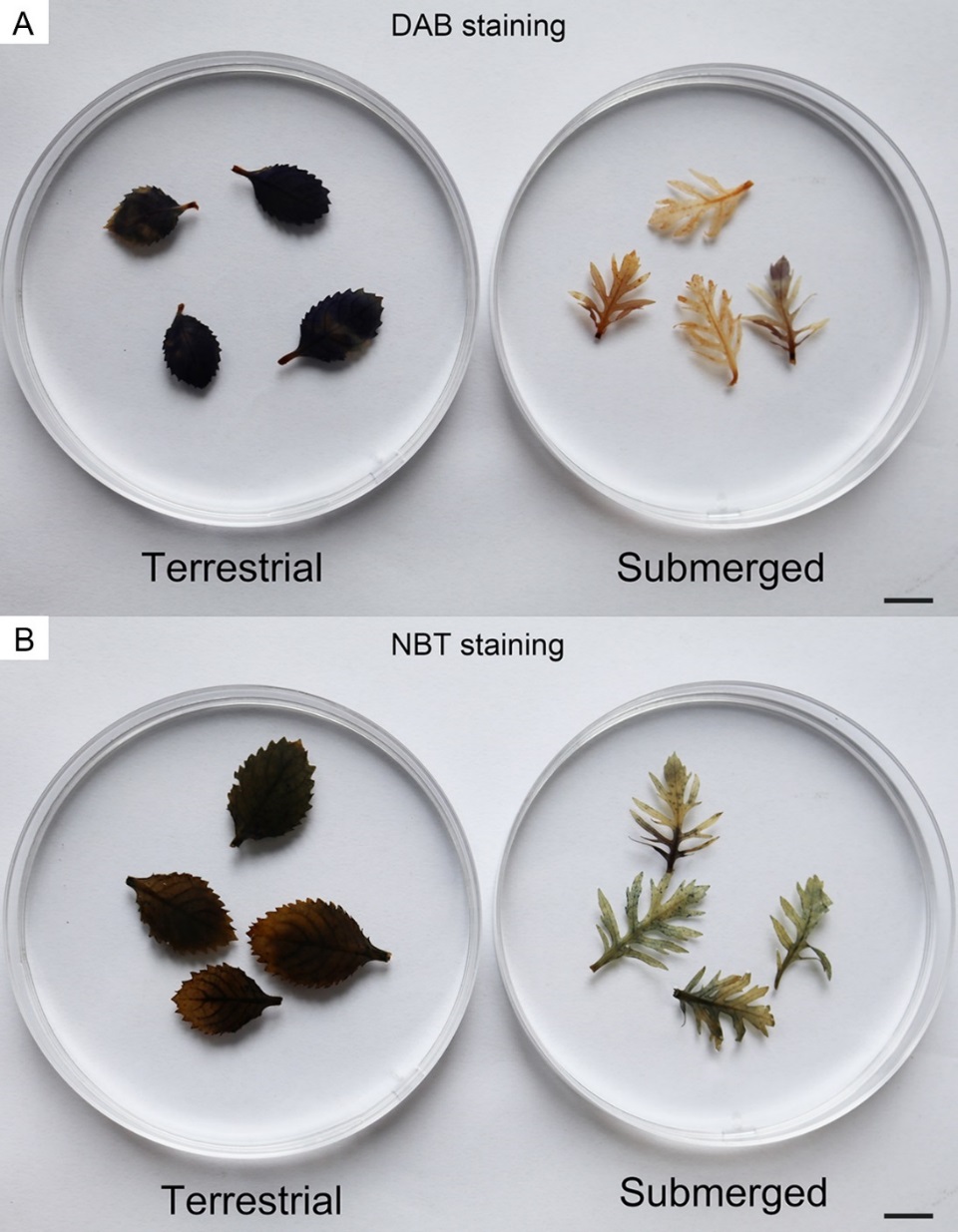

**Figure S7.** ROS detection in terrestrial or submerged leaves. DAB staining (A) and NBT staining (B). Photos show four representative P6 stage leaves from the twenty independent plants grown in each condition. Bars = 1 cm.

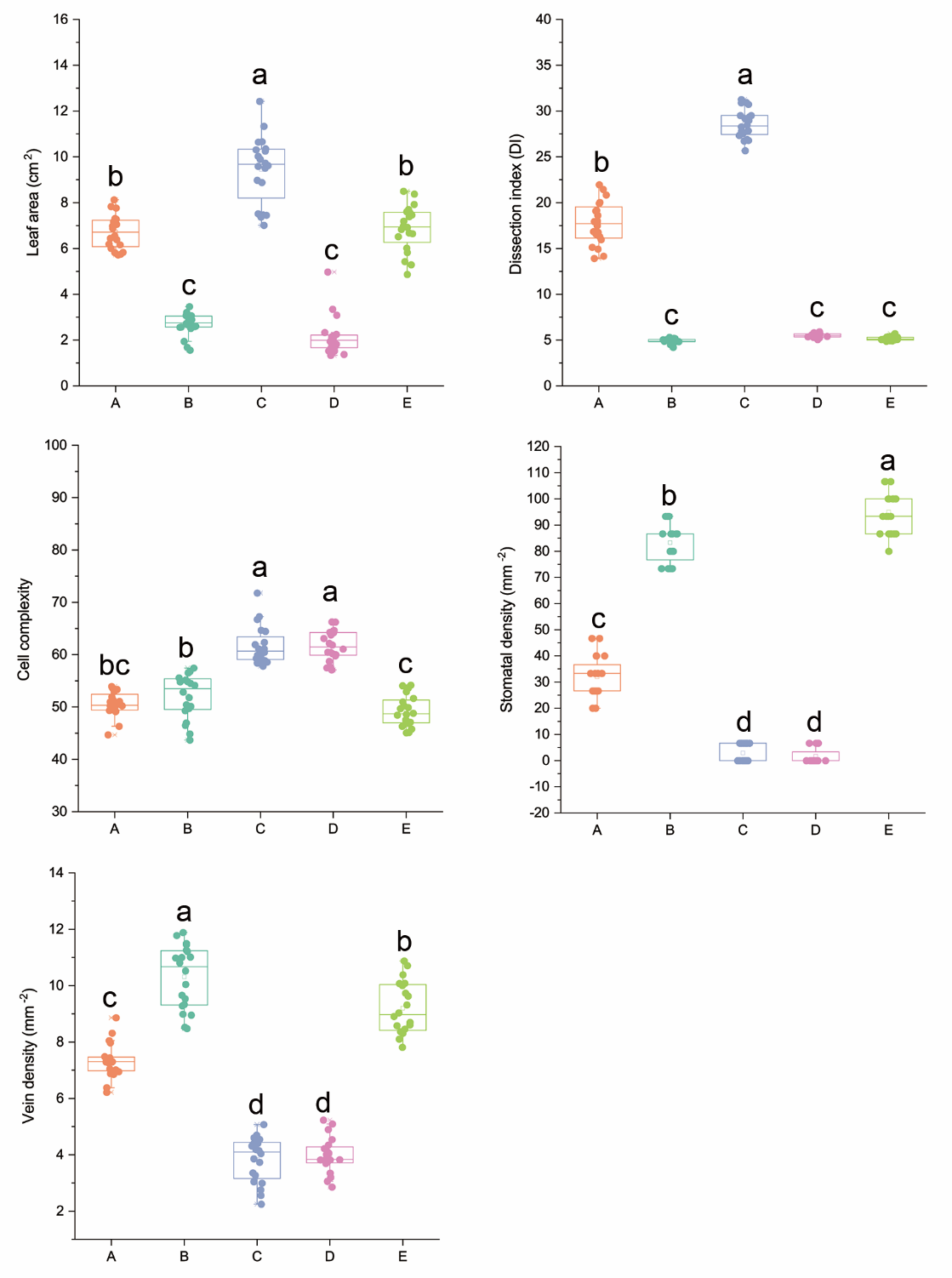

**Figure S8.** Morphological parameters of *H. difformis* in different groups. (A) terrestrial plants grown at 23°C, 60% RH, 60 μmol m^-2^ s^-1^ illumination; (B) terrestrial plants grown at 18°C, 60% RH, 60 μmol m^-2^ s^-1^ illumination; (C) submerged plants grown at 23°C, 60 μmol m^-2^ s^-1^ illumination; (D) submerged plants grown at 23°C, 10 μmol m^-2^ s^-1^ illumination; (E) terrestrial plants grown at 23°C, 30% RH, 60 μmol m^-2^ s^-1^ illumination). Data are from twenty biological replicates for each condition, and lowercase letters indicate significant differences determined by Tukey’s test of one-way ANOVA (*P* < 0.05).

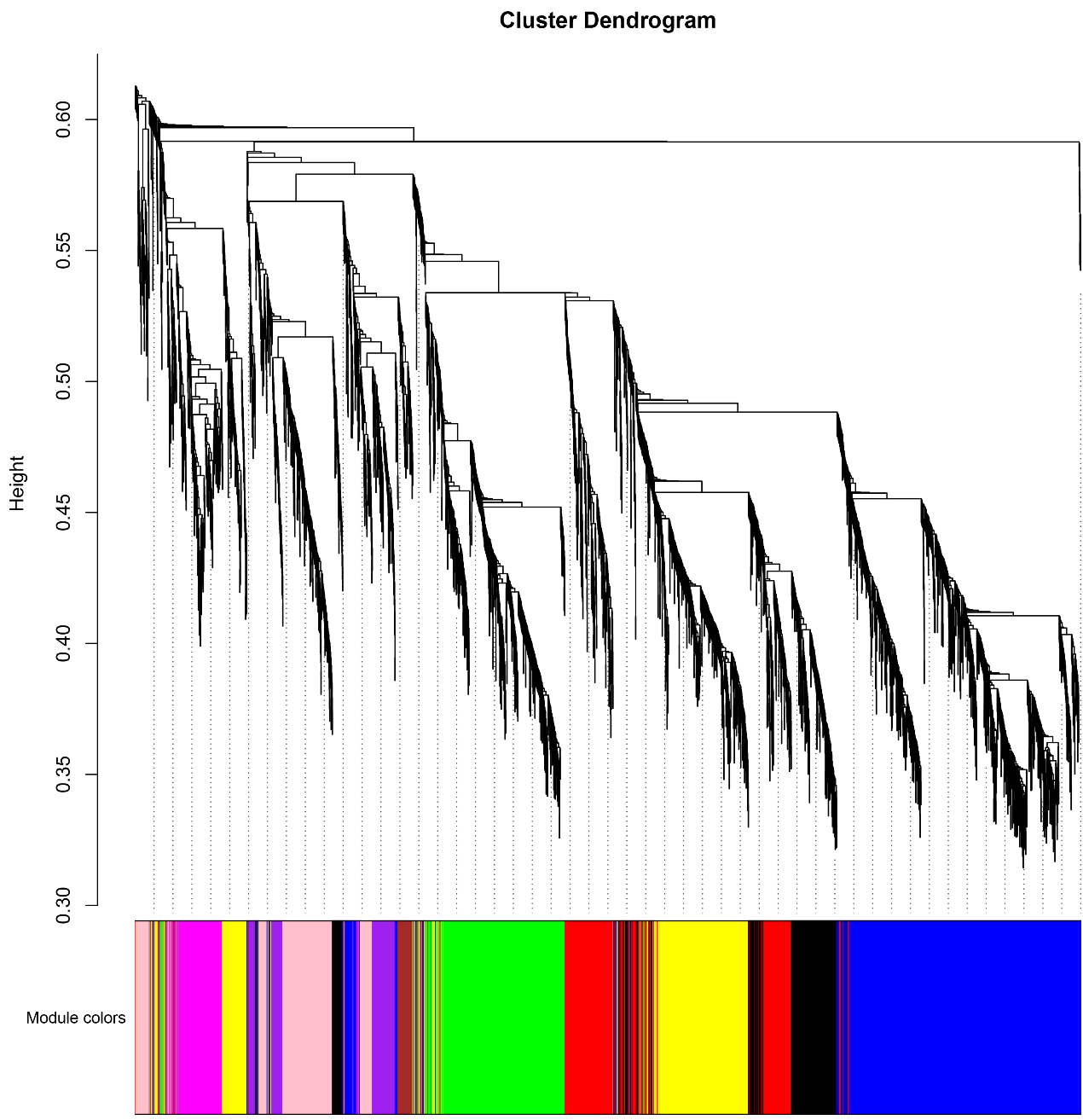

**Figure S9.** Gene dendrogram and corresponding module colors. The clustering was based on expression levels of filtered DEGs.

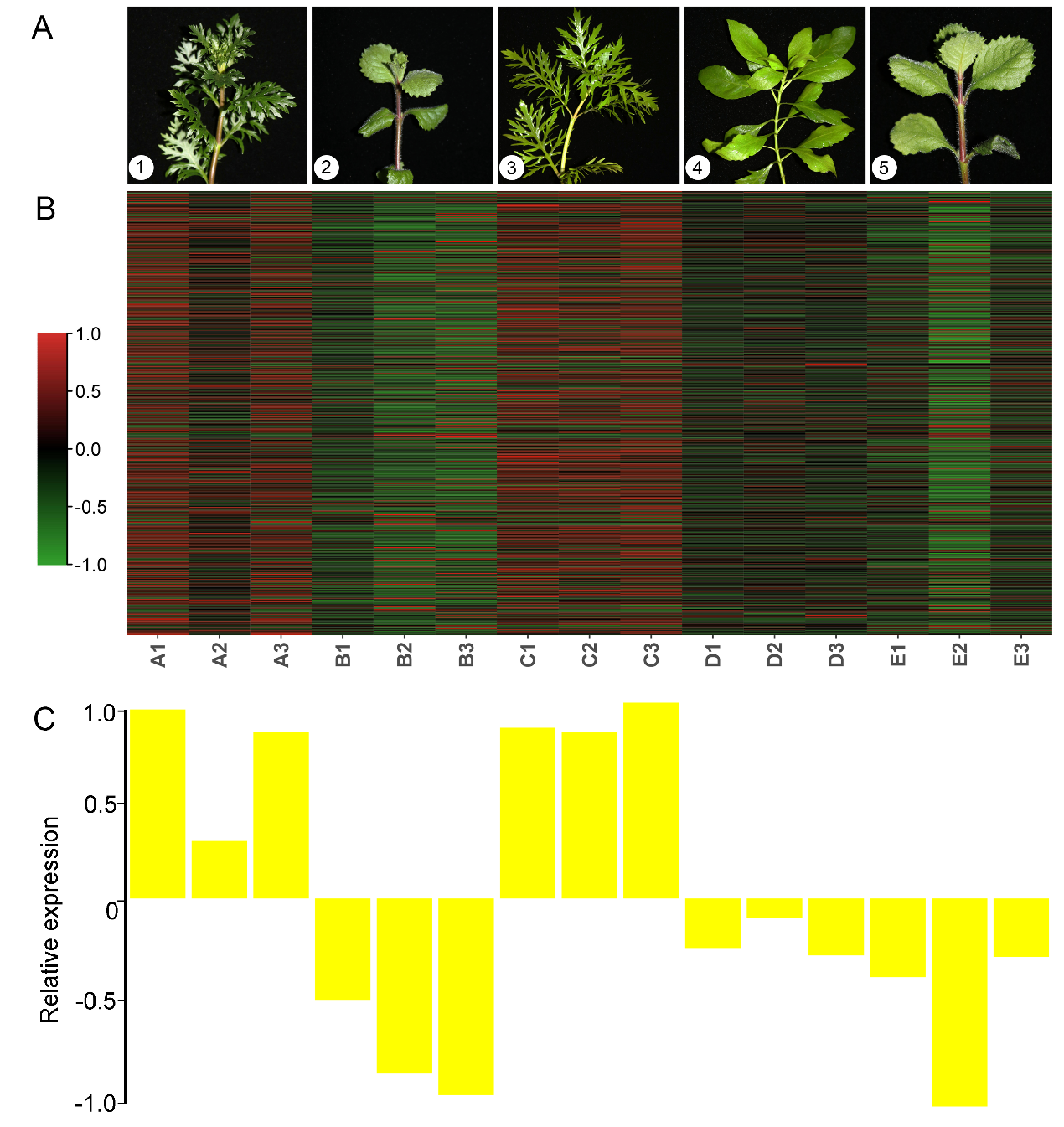

**Figure S10.** Co-expression network and phenotypic analysis in the yellow module.

(A) Phenotypes of *H. difformis* grown in different conditions (1: terrestrial plants grown at 23°C, 60% RH, 60 μmol m^-2^ s^-1^ illumination; 2: terrestrial plants grown at 18°C, 60% RH, 60 μmol m^-2^ s^-1^ illumination; 3: submerged plants grown at 23°C, 60 μmol m^-2^ s^-1^ illumination, 4: submerged plants grown at 23°C, 10 μmol m^-2^ s^-1^ illumination, 5: terrestrial plants grown at 23°C, 30% RH, 60 μmol m^-2^ s^-1^ illumination).

(B) Heatmaps of genes belonging to the yellow module. Red indicates upregulated genes and green indicates downregulated genes.
